# Sustained ERK signaling couples the injury response to organizer formation during *Hydra* head regeneration

**DOI:** 10.64898/2025.12.27.696609

**Authors:** I.Y. Juanico, A.W. Stockinger, A.K. Virgen, N. Srisrimal, S.E. Campos, C.E. Juliano

## Abstract

Regenerative abilities vary widely across animals, even among closely related species, and a central challenge is to compare their gene regulatory networks (**GRNs**) to determine which components and regulatory connections are conserved or divergent. *Hydra* provides a powerful system for this work: as a cnidarian, which is the sister group to bilaterians, it offers access to deeply conserved regenerative mechanisms, and species within the genus differ in regenerative phenotypes. Here, we investigate the function of extracellular signal-regulated kinase (**ERK**) signaling in driving head regeneration in *Hydra*. ERK is a conserved injury-responsive pathway implicated in regeneration across animals, including in the activation of Wnt signaling, which is necessary and sufficient to establish the *Hydra* head organizer, a Wnt signaling center that patterns the main body axis. We found that in *Hydra vulgaris*, ERK activity extends from the generic injury phase into the period when the Wnt-based head organizer forms. However, unlike in some other regenerating species, transient ERK inhibition delayed but did not block regeneration, consistent with *Hydra’s* ability to regenerate without a proliferation-dependent blastema. ERK inhibition in *H. vulgaris* disrupted the transcriptional activation of multiple Wnt pathway components and restricted their expression domains, providing a likely explanation for the delayed onset of head organizer formation. Comparative experiments in *Hydra oligactis*, which regenerates its head approximately 24 hours more slowly than *H. vulgaris*, revealed reduced persistence of ERK signaling and lower sensitivity to ERK inhibition. These features suggest that while injury-induced ERK activity is conserved, the mechanisms that establish the Wnt signaling center during regeneration diverge between the two species. Together, our findings identify sustained ERK activity as a critical regulator of Wnt pathway activation during *Hydra* head regeneration and indicate that evolutionary shifts in the ERK-Wnt regulatory architecture contribute to species-specific differences in regenerative phenotypes.

## INTRODUCTION

Developmental processes and the molecular response to injury are distinct biological programs, each conserved across animals. Regeneration requires linking these two programs, yet the nature of this link varies widely across species, suggesting substantial evolutionary flexibility in how injury signals trigger developmental patterning. Some animals such as planarians^1^ and *Nematostella*^2^ are capable of whole-body regeneration, while others are able to regenerate specific structures, such as the axolotl limb^3^ and the zebrafish caudal fin.^4^ Even closely related species can differ substantially in their regenerative capacity. For example, the spiny mouse *Acomys* exhibits exceptional regeneration of skin and muscle^5,6^ in comparison to the more limited healing response of the common laboratory mouse *Mus musculus.* In addition, planarian species also differ in their regenerative abilities, driven in part by species-specific differences in Wnt signaling deployment.^7,8^ This variability of regenerative ability, even among closely related species, suggests that conserved components are wired differently into gene regulatory networks (**GRNs**) to produce distinct regenerative outcomes.^9^ The GRNs that coordinate the injury response to trigger the regeneration of missing body parts remain incompletely defined in any species. Defining these networks in highly regenerative animals will uncover how conserved developmental pathways are activated to restore missing tissue and how this ability is lost and gained repeatedly during evolution.^10,11^

The ERK signaling pathway is a MAPK cascade that forms a core component of the conserved injury response in both regenerative and non-regenerative animals and is essential in many regenerative contexts. For example, ear punch injury activates ERK signaling in both *Mus musculus* and *Acomys,* yet only *Acomys* fully regenerates.^6^ Experimentally boosting ERK signaling activity in *Mus musculus* improves the regenerative outcome.^6^ ERK signaling is also required for regeneration of the caudal fin and osteoblasts in zebrafish^12,13^ and limb regeneration in the American cockroach,^14^ *Xenopus*,^15^ and salamanders.^16^ Whole-body regeneration in planaria,^17,18^ acoels,^19^ and *Nematostella vectensis*^20^ also require ERK signaling. Downstream targets of ERK include conserved immediate early genes (**IEGs**) such as bZIP and EGR family TFs.^21^ Although these factors function broadly in wound healing, they are also implicated in activating Wnt signaling in regenerative contexts. In *acoels*, for example, injury-induced ERK is required for *egr* transcription, which in turn transcriptionally activates Wnt pathway genes.^19,22^ In planarians, ERK signaling is required for the activation of Wnt signaling components.^17^ In *Drosophila*, the bZIP transcription factor (**TF**) complex AP-1 activates Wnt signaling during regeneration of the wing disc, and this activation depends on JNK signaling, another MAPK pathway.^23,24^ Together, these studies suggest a conserved molecular toolkit in which MAPK signaling promotes Wnt pathway activation through IEG TFs.^9^ However, the precise mechanisms of these interactions, and how they have been maintained or modified across evolution, remain unclear.

*Hydra* provides a powerful model for understanding how the injury response shapes regenerative outcomes, as well as how these processes evolve. *Hydra* is a freshwater cnidarian polyp with a simple body plan organized around an oral-aboral axis, with a head at one end and a foot at the other (Fig. 1A). The most commonly studied *Hydra* species, *Hydra vulgaris*, reliably regenerates both its head and foot after bisection of the body column, a region that contains proliferative stem cells.^25^ The body structure is formed by two epithelial layers, the ectoderm and the endoderm, and the epithelial cells in the body column are constitutively proliferative. These cells retain the capacity to regenerate either head or foot structures in response to injury, enabling robust and repeatable regeneration.^26^ Following injury in the form of bisection perpendicular to the oral-aboral axis, ERK signaling is rapidly activated at both wound sites and remains elevated for at least 6 hours.^27^ Furthermore, inhibiting ERK signaling with the drug U0126, which targets MEK, reduces head and foot regeneration rates after bisection.^27,28^ However, the duration of the ERK signaling injury response in *H. vulgaris* and the timing of its action during regeneration remains unclear. A key advantage of *Hydra* for studying how ERK signaling activates developmental pathways is that unlike many other regeneration models, an initial burst of proliferation is not required for regeneration.^29^ This distinction uncouples ERK’s roles in promoting proliferation from its role in directing transcription of patterning gnes,^26,30,31^ allowing us to focus specifically on ERK-dependent patterning programs in *Hydra*.

**Fig 1.**
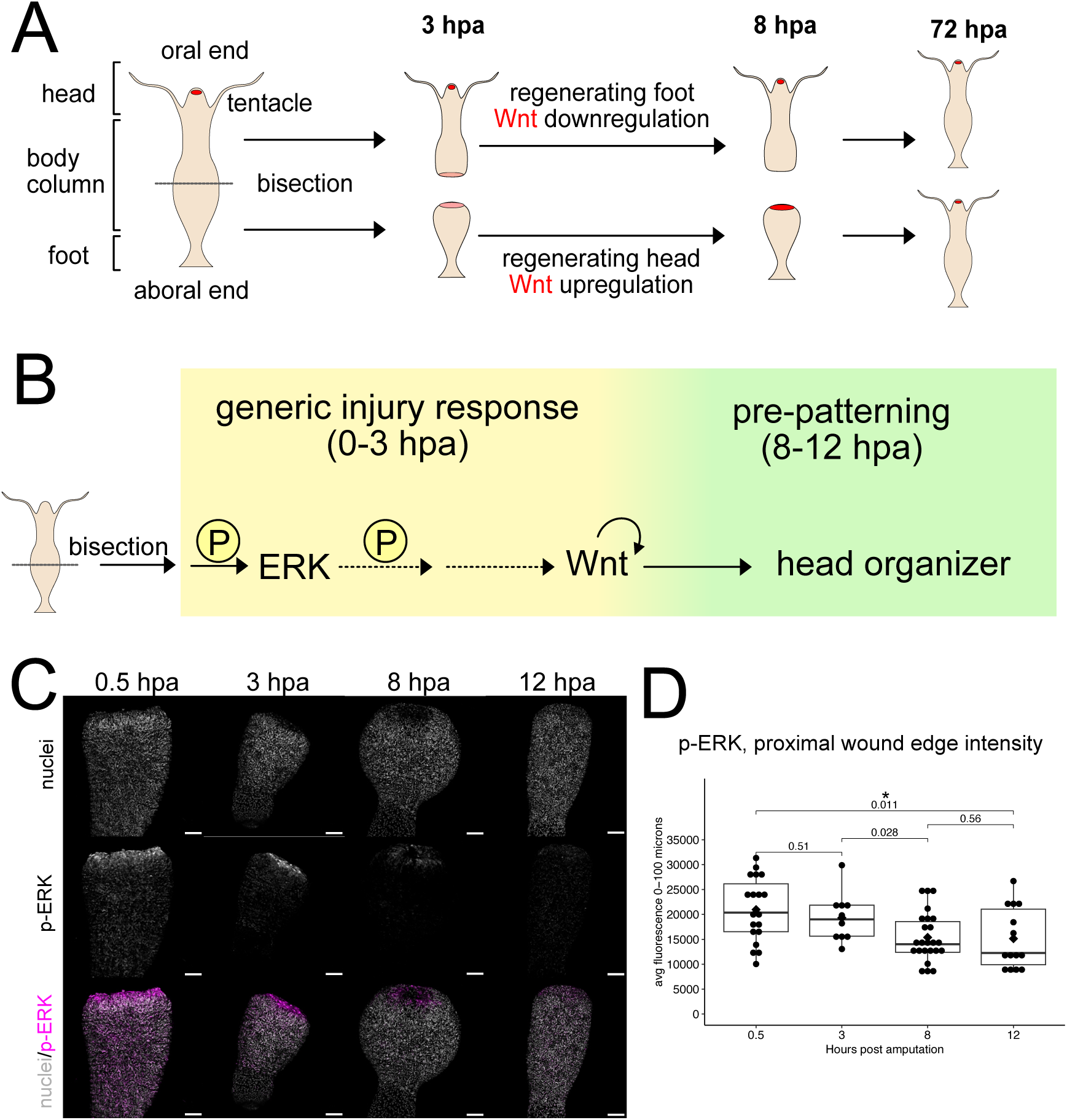
ERK activation dynamics and a proposed role for ERK in driving Wnt signaling during *Hydra vulgaris* head regeneration. (A) Model of *H. vulgaris* head and foot regeneration showing the spatial and temporal dynamics of Wnt transcript levels (red).^36^ (B) Hypothesized model for how injury-induced ERK signaling activates Wnt transcription during head regeneration in *H. vulgaris*.^27,36^ Solid arrows indicate regulatory relationships supported by published studies, while dotted arrows represent proposed connections suggested by current evidence but not yet fully established. (C) Immunofluorescence of regenerating *H. vulgaris* heads using anti-phospho-ERK (p-ERK) together with Hoechst to label nuclei shows detectable expression at the wound edge through 12 hours post amputation (hpa). Scale bar =100µm. (D) Quantification of the p-ERK signal from panel C, with each point representing an individual animal (0.5 hpa n=19, 3 hpa n=10, 8 hpa n=24, 12 hpa n=14). Fluorescence intensity was measured using a plot profile analysis. Statistical comparisons were performed using the Wilcoxon rank-sum test.

The oral-aboral axis in *Hydra* is established by a Wnt signaling center at the oral end that is necessary and sufficient for head patterning.^32–35^ However, Wnt signaling is also activated during the generic injury phase, including at non-regenerative wounds and when regenerating head and foot tissues are transcriptionally indistinguishable (Fig. 1A).^34,36^ As regeneration progresses and the tissue acquires structure-specific identity, Wnt continues to be transcriptionally upregulated in the regenerating head and downregulated in the regenerating foot (Fig. 1A).^34,36^ This structure-specific phase of regeneration, in which the tissue identity is established before morphological features appear, occurs between 8 and 12 hpa and is referred to as the pre-patterning phase (Fig. 1B). There is evidence that ERK signaling is required for the activation of Wnt signaling and head regeneration,^27^ but it remains unclear whether ERK is required during the generic injury phase, the head pre-patterning phase, or both. Being able to define the GRNs underlying these two distinct phases of Wnt activation is critical for understanding how injury signals are translated into precise patterning outcomes.

Comparative studies are a powerful tool for dissecting biological mechanisms. As the sister group to bilaterians, cnidarians such as *Hydra* are well suited for identifying both conserved and divergent principles of regeneration.^37^ In addition, while broad comparisons across distant clades are clearly valuable, comparisons between closely related species allow the underlying regulatory logic to be resolved more precisely and expose specific GRN changes that drive differences in regeneration. *Hydra oligactis* provides such a comparison to *H. vulgaris*; although the two species are separated by only ∼30 million years of evolution^38,39^ and share a similar body plan, *H. oligactis* is less regenerative. Compared to *H. vulgaris*, *H. oligactis* displays a delay in head regeneration and has significantly reduced rates of foot regeneration.^40,41^ Our recent study leveraged these phenotypic differences to show that a reduction of injury-induced Wnt signaling, due to low transcription of Wnt signaling genes, contributes to the regeneration phenotype differences observed in *H. oligactis* as compared to *H. vulgaris*.^41^ However, the basis of this reduced transcriptional response during injury remains unclear. Comparing upstream signaling pathways such as ERK between these two species will clarify the architecture of the regeneration GRN and reveal how its regulation has evolved to generate phenotypic divergence.

In this study, we further elucidate the GRN that drives regeneration in *Hydra* by defining the injury-induced dynamics and timing of ERK signaling function during *H. vulgaris* head regeneration. We find that injury-induced ERK signaling extends into the pre-patterning phase and maintains functional roles beyond the initial injury response. ERK activity is required both for the injury-induced transcriptional activation of a subset of Wnt genes and for the expansion of the injury domain into surrounding tissue. We also show that ERK dynamics differ between *H. oligactis* and *H. vulgaris*; specifically, ERK signaling attenuates more rapidly in *H. oligactis*, and head regeneration in this species is correspondingly less sensitive to ERK inhibition. These observations suggest that ERK signaling is decoupled from the rapid activation of Wnt signaling in *H. oligactis*, contributing to its slower overall timeline of head regeneration. Together, these findings reveal key features of ERK input into the *H. vulgaris* regeneration GRN, clarify ERK’s role in directing early patterning, and illustrate how differences in the persistence of ERK activity may underlie evolutionary variation in regenerative capacity.

## RESULTS

### ERK signaling is required beyond the generic injury response to enable rapid head regeneration in Hydra vulgaris

A previous study showed that continuous inhibition of ERK signaling with the MEK inhibitor U0126 had inhibitory effects on regeneration and moderately reduced expression levels of two Wnt ligand genes, *wnt3* and *wnt9/10c*, in multiple injury contexts.^31^ These findings support the prevailing hypothesis that injury-induced ERK signaling activates the generic transcription of Wnt signaling genes at all wound sites and that this activation is then selectively sustained at oral-facing wounds through a feedforward loop in the regenerating head to establish the oral organizer (Fig. 1A and B).^27,36^ However, because previous studies were done with continuous ERK inhibition,^27^ it remained unclear if ERK signaling is only required to activate Wnt transcription during the generic injury phase, which resolves between 3 and 8 hpa, or if this requirement continues into the pre-patterning phase, which occurs between 8 and 12 hpa (Fig. 1B). To address this gap, we first tested the timeframe of ERK signaling activation during *H. vulgaris* head regeneration. The previous study reported a strong p-ERK signal by Western blot through 6 hpa, but the experiment was not extended through the pre-patterning phase.^27^ We therefore used immunofluorescence to measure p-ERK signal at the wound edge and in the adjacent tissue at multiple timepoints through 12 hpa to reveal the dynamics through the pre-patterning phase of regeneration. Similar to published results,^27^ we observed a strong signal at the proximal wound edge (the first 100µm from the edge of the tissue) in response to bisection. This signal remained elevated through 8 hpa and then decreased, reaching significantly lower levels than at 0.5 by 12 hpa (Fig. 1C and D). Notably, some animals still demonstrated visible p-ERK signal at the wound edge even at 12 hpa (Supp. Fig. 1). Similarly, injury-induced p-ERK signal distal to the wound (200-400µm from the wound edge) was elevated through 8 hpa and then decreased significantly compared to 0.5 hpa by 12h pa (Supp. Fig. 2A). Altogether, we found that injury induced an increase of p-ERK at the wound site that is propagated into the surrounding tissue and remains elevated until at least 12 hpa, well into the pre-patterning phase of *H. vulgaris* regeneration.

We next aimed to delineate the requirement of ERK signaling during the generic injury phase versus the pre-patterning phase of head regeneration (Fig. 1B). To accomplish this, we used two ERK inhibitors with different mechanisms of action, U0126 and BVD-523, to control for off-target effects. U0126 is an inhibitor of MEK, the kinase that directly phosphorylates ERK, that has been previously validated to reduce p-ERK levels in *H. vulgaris* during regeneration.^27^ Consistent with previous studies, bisected animals treated with U0126 showed a significantly reduced p-ERK signal, confirming the effectiveness of the inhibitor in our experiments (Supp. Fig. 3A, C, and D).^27^ BVD-523 is an ATP-competitive ERK inhibitor that paradoxically increases detectable levels of p-ERK levels in *H. vulgaris* (Supp. Fig. 3D and Supp. Fig. 4A), consistent with reports in other systems.^42–44^ This effect occurs because BVD-523 inhibits the catalytic activity of ERK, preventing phosphorylation of downstream targets, but it does not block MEK from phosphorylating ERK.^45^ As a result, catalytically inactive p-ERK accumulates. Thus, the observed increase in p-ERK following BVD-523 treatment indicates effective inhibition of ERK signaling in *H. vulgaris*. For regeneration assays, we selected a concentration of 7.5 µM BVD-523, the highest concentration that did not cause significant homeostatic disruption, such as tentacle shortening or loss, after a 24-hour incubation.

To test whether ERK signaling is required only during the generic injury response or also during the pre-patterning phase of head regeneration, we transiently treated *H. vulgaris* with either 25 μM U0126, 7.5 μM BVD-523, or DMSO, the vehicle control, for the first 3, 8, or 12 hpa, then washed the animals back into *Hydra* medium (HM). In all experiments, treatment began one hour before bisection, which was perpendicular to the oral-aboral axis and midway between the head and foot (i.e., 50% bisection, see Fig. 1A,B schematic). Animals were imaged and evaluated every 24 hours, and the emergence of two or more tentacle buds was scored as head regeneration. Inhibition of ERK signaling during the first 3 hpa had no effect on the rate of head regeneration, and extending inhibition to 8 hpa caused only a mild head regeneration delay (Fig. 2A). However, inhibition for 12 hpa, which is well beyond the generic injury phase and into the pre-patterning phase, produced a clear and reproducible ∼24-hour delay with both inhibitors. All ERK inhibited regenerating animals successfully closed their wounds and were morphologically normal (Fig. 2B). The head regeneration delay was most apparent at 48 hpa, when there are significantly lower numbers of tentacles in ERK inhibited regenerating animals as compared to the DMSO vehicle control (Fig. 2C).

**Fig. 2.**
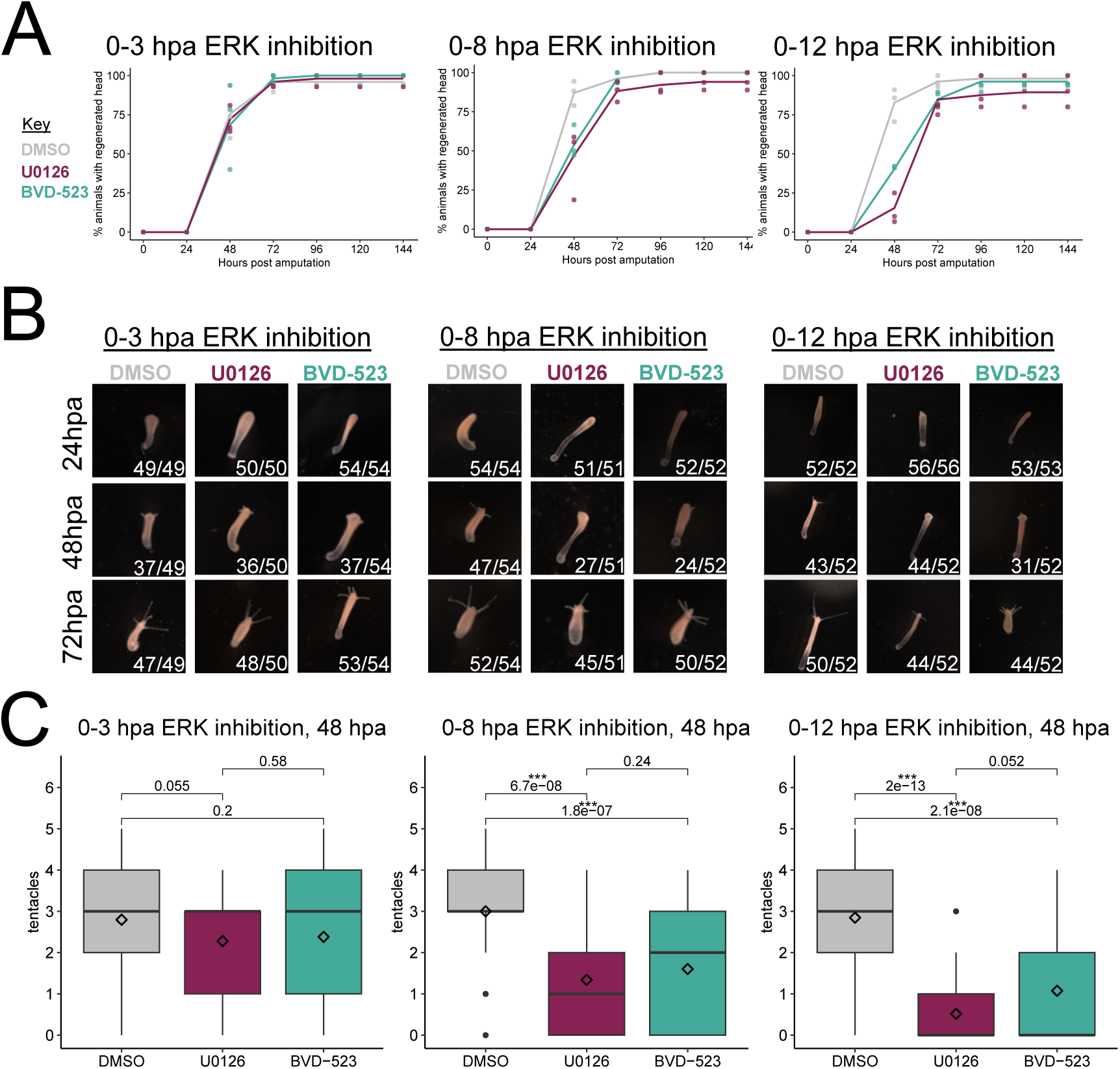
Transient ERK signaling activity is required for rapid head regeneration in *H. vulgaris*. (A) Head regeneration rates for *H. vulgaris* treated with ERK inhibitors (U0126, 25µM, burgundy; BVD-523, 7.5µM, teal) or a vehicle control (DMSO, gray) for 0-3 hpa, 0-8 hpa, or 0-12 hpa, then transferred to DMSO until 12 hpa. After 12 hpa, all animals were washed with *Hydra* medium for the remainder of the experiment. Each point represents a biological replicate. Sample sizes were as follows: 0-12 hpa treatment (DMSO n=52, U0126 n=56, BVD-523 n=53), 0-8 hpa treatment (DMSO n=54, U0126 n=51, BVD-523 n=52), and 0-3 hpa treatment (DMSO n=49, U0126 n=50, BVD-523, n=54). (B) Representative images of *H. vulgaris* head regeneration at 24, 48, and 72 hpa from treatments shown in panel A. (C) Quantification of tentacle number at 48 hpa for control (DMSO, gray) and ERK inhibitor-treated (U0126, burgundy; BVD-523 teal) animals from experiments shown in panel A. Tentacles from all three biological replicates were pooled for analysis. The diamond represents the mean. Statistical comparisons were performed using the Wilcoxon rank-sum test.

One explanation for why head regeneration is delayed, rather than blocked, is that p-ERK signal recovers after removal of the inhibitor. To test this, we performed Western blot analysis to measure p-ERK levels one hour after the 0-3 hpa, 0-8 hpa, and 0-12 hpa U0126 experiments (Fig. 3A). We observed a significant recovery of the p-ERK signal one hour after the 0-3 hpa treatment, which is consistent with the absence of a regeneration defect under these conditions (Figs. 2A and 3B, Supp. Figs. 4B, C). By contrast, recovery was reduced after the 0–8 hpa washout and was nearly absent after the 0–12 hpa washout, correlating with progressively slower regeneration rates (Figs. 2A and 3B, Supp. Fig. 4B, C). These results indicate that the function of ERK signaling during the generic injury phase extends into the pre-patterning phase of head regeneration. Restricting U0126 exposure to 12–24 hpa also produces a ∼24-hour delay in head regeneration (Supp. Figs. 3E and F), indicating that p-ERK activity remains functionally important even beyond 12 hpa, consistent with the basal p-ERK signal observed in uninjured tissue (Supp. Fig. 5A).

**Fig. 3.**
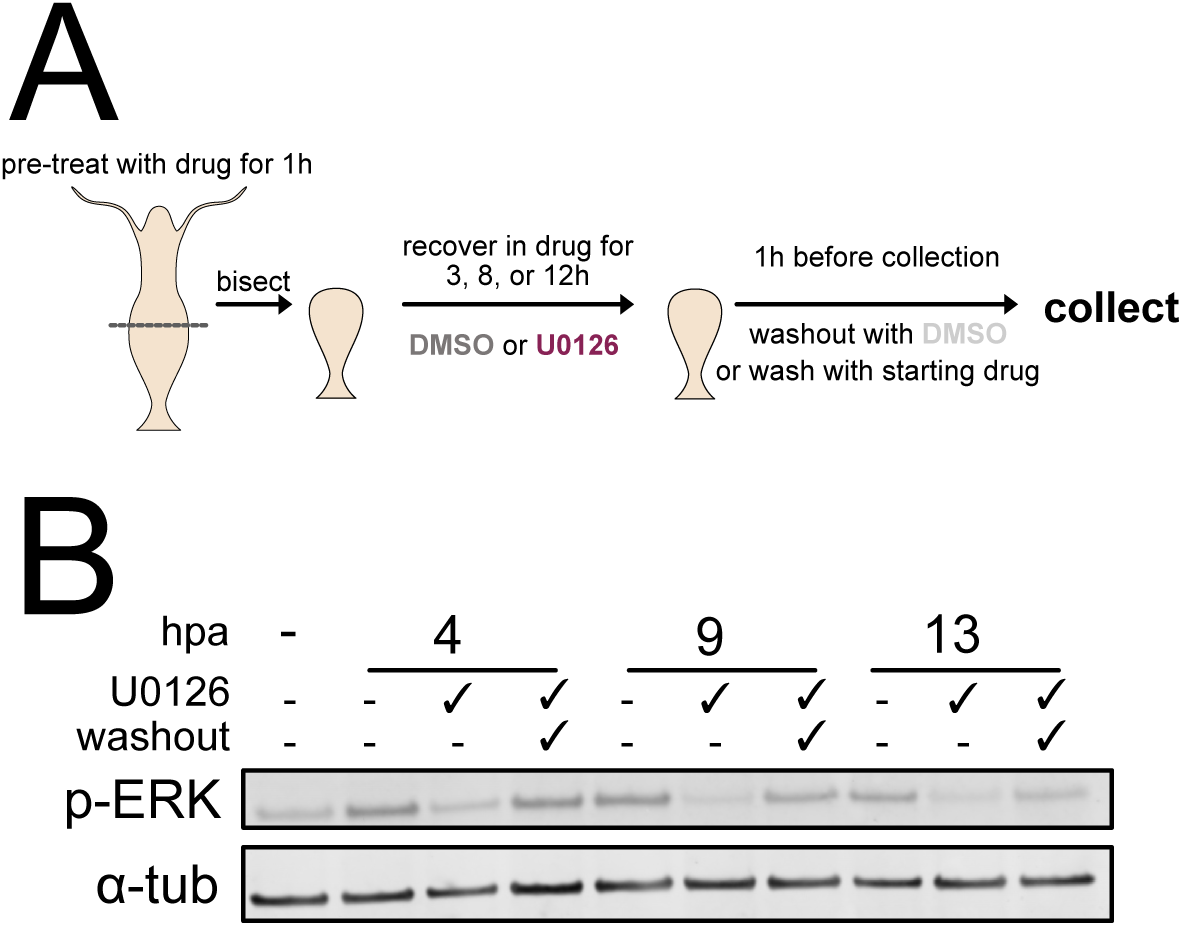
ERK signaling activity partially recovers after transient inhibition. (A) Schematic of the experiment used to assess the recovery of p-ERK signal after wash out of U0126. (B) Western Blot showing p-ERK and α-tubulin levels in regenerating *H. vulgaris* heads. Animals not treated with U0126 received DMSO as a vehicle control. “Washout” indicates that animals were washed into *Hydra* Medium 1 hour before tissue collection. The first lane of the blot shows whole animals with no injury or treatment, indicating basal levels of p-ERK.

Cell proliferation is not strictly required for *Hydra* head regeneration, although inhibiting it can delay the process.^29^ Furthermore, ERK signaling has a well-established role in promoting cell proliferation, including in many regenerative contexts. ^6,14,17,18,46,47^ We therefore next tested whether the regeneration delay observed under ERK inhibition might result from reduced cell proliferation. To accomplish this, we measured proliferation using phospho-histone-3 (pH3) and Hoechst staining on whole-mount regenerating heads treated with U0126, BVD-523, or DMSO for 0-3 hpa, 0-8 hpa, and 0-12 hpa and found no significant difference in the number of pH3-positive nuclei (Fig. 4 and Supp. Fig. 6). These data show that ERK inhibition does not significantly affect cell proliferation in regenerating heads during the transition from injury to the pre-patterning phase, consistent with prior reports that early head regeneration in *H. vulgaris* does not involve or require increased cell proliferation.^26,31^

**Fig. 4.**
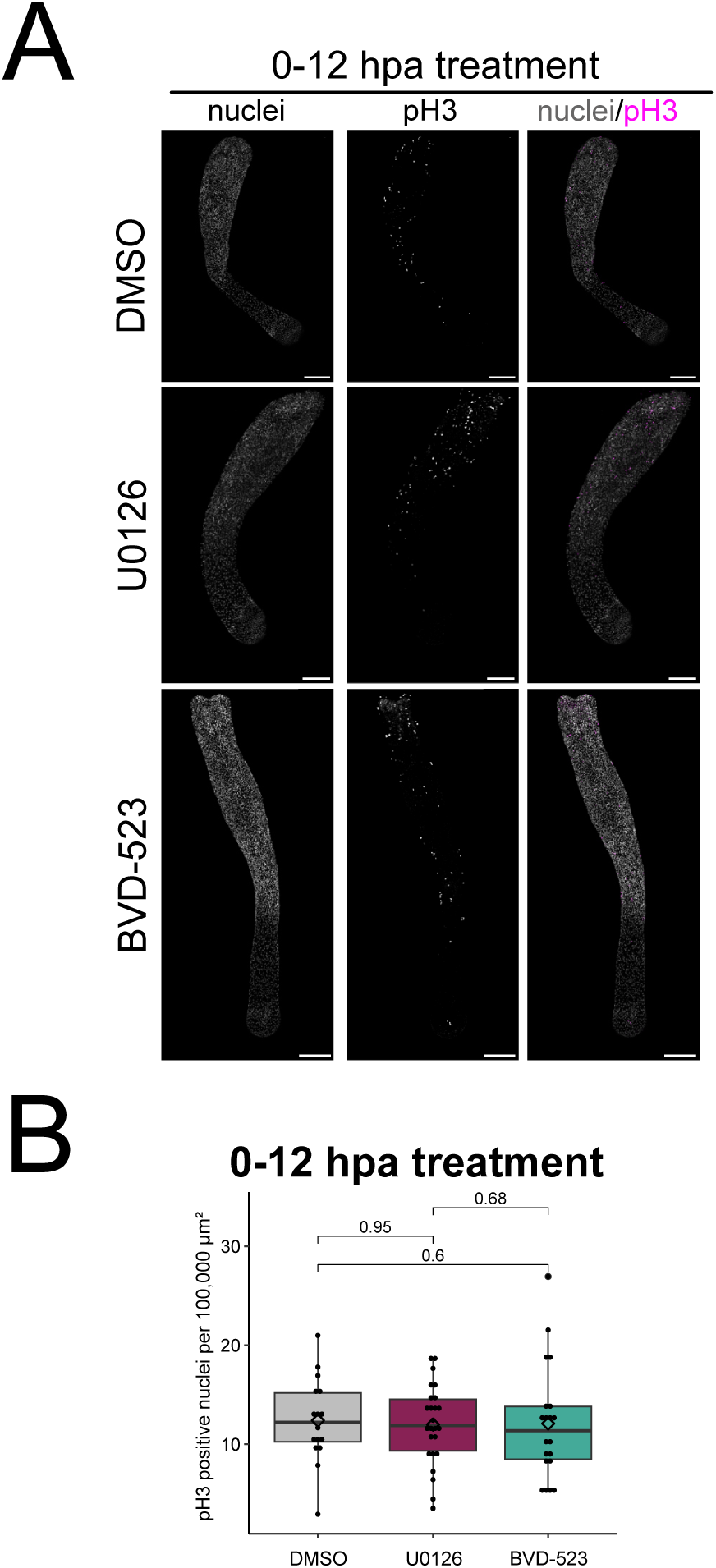
ERK inhibition does not affect proliferation during early head regeneration in *H. vulgaris*. (A) Representative images of *H. vulgaris* treated with ERK inhibitors (U0126 or BVD-523) or DMSO vehicle control for the first 12 hours of regeneration stained with pH3 antibody to identify mitotic cells and counterstained with Hoechst (nuclei). Scale bar = 100µm. (B) Quantification of pH3-positive nuclei normalized by tissue area for the experiment described in panel A (DMSO n=16, U0126 n=25, BVD-523 n=20). Statistical comparisons were performed using the Wilcoxon rank-sum test.

### ERK activity is required for the injury-induced transcription of a subset of Wnt signaling genes

ERK signaling inhibition for the first 12 hours after bisection delayed head regeneration (Fig. 2A), suggesting that ERK activity is required for the timely transcriptional activation of head patterning genes. In support of this, a previous study reported subtle transcriptional changes in two Wnt ligand genes during regeneration through 6 hpa using qPCR.^27^ To test this more comprehensively and to extend the timeline to include the pre-patterning phase, we performed RNA-seq on regenerating *Hydra* head tissue at 0, 3, 8, and 12 hpa treated with U0126 or the vehicle control DMSO (Fig. 5A), then performed differential gene expression (**DGE**) analysis (Supp. Table 1).

**Fig. 5.**
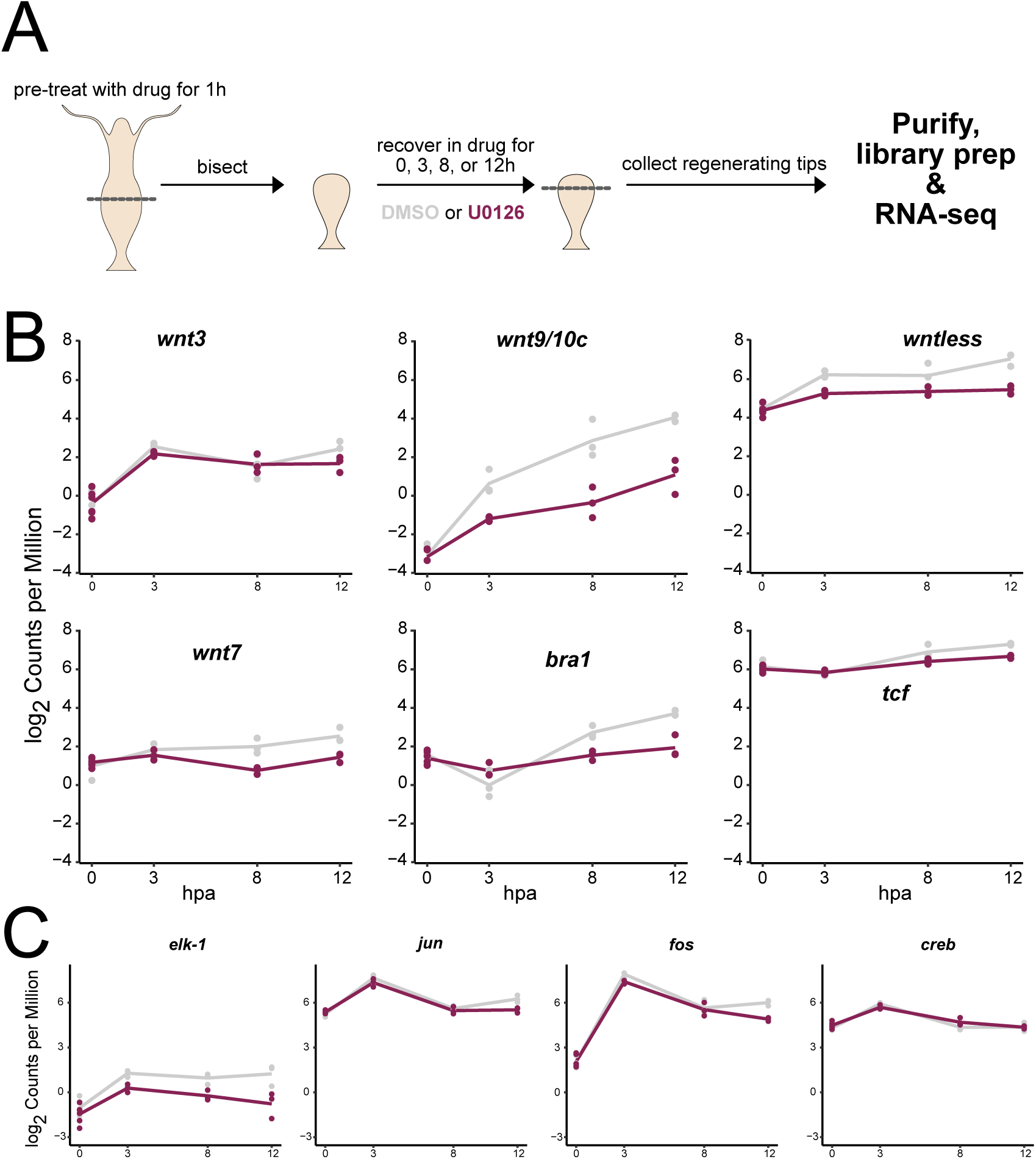
ERK signaling activity is required for the transcription of selected Wnt signaling genes, but not bZIP transcription factor genes in *H. vulgaris.* (A) Schematic of RNA-seq experiment to test the effect of ERK signaling inhibition on gene expression. (B) Transcript levels of *wnt3, wnt9/10c, wntless, wnt7*, *bra1,* and *tcf* in animals treated with U0126 (burgundy) or DMSO vehicle control (gray) collected at 0, 3, 8, and 12 hpa. Each point represents a biological replicate. (C) Transcript levels of *elk-1* (G004172), *jun, fos,* and *creb* in animals treated with U0126 (burgundy) and DMSO vehicle control (gray), collected at 0, 3, 8, and 12 hpa. Each point represents a biological replicate.

Homeostatic animals exhibit a basal p-ERK signal, indicating that ERK signaling has roles in this context as well as regeneration (Supp. Fig. 5). Therefore, to distinguish regeneration-specific effects, we also performed RNA-seq and DGE analysis on uninjured animals treated with U0126 or DMSO for 12 hours. We then compared the differentially expressed genes (**DEGs**) from ERK-inhibited homeostatic animals with those from ERK-inhibited regenerating animals (Supp. Table 2). We found very little overlap between these DEGs (Supp. Fig. 7), indicating that ERK signaling has different functions in homeostasis as compared to regeneration. While the homeostatic function of ERK signaling is not the primary focus of our study, by intersecting genes that are differentially expressed following ERK inhibition in homeostatic animals with our single-cell expression atlas,^48^ we can report that the effect is larger in the ectodermal and endodermal epithelial lineages (Supp. Fig. 8).

We next focused on the transcriptional changes caused by ERK inhibition during head regeneration. Differentially expressed genes (**DEGs**) in response to ERK inhibition in regenerating animals are largely expressed in the ectodermal and endodermal epithelial lineages in our whole animal scRNA-seq atlas, suggesting the effect is strongest in these lineages during regeneration (Supp. Fig. 9). We next used the regression-based tool maSigPro to identify DEGs with similar temporal expression dynamics, which infers co-regulation or shared biological function (Supp. Fig. 10A).^49^ Clusters 5 and 9 included genes downregulated by ERK inhibition. Notably, the Wnt pathway genes *wntless* (G013231) and *wnt7* (G023882) were found in Cluster 5, while *wnt9/10c* (G011290), *tcf* (G007593), and *brachyury1* (G006413), a known Wnt target,^50^ appear in Cluster 9 (Supp. Fig. 10A, Supp. Table 3, Fig. 5B). Although only *wntless* and *brachyury1* met our significance threshold for downregulation, the overall trend across the dataset indicates reduced Wnt pathway activity, reinforced by the significant downregulation of the Wnt target *brachyury1*. This is consistent with regulation of the Wnt signaling pathway being enriched in our GO analysis 12 hpa (Supp. Fig. 11). Interestingly, *wnt3* (G010730) did not fall into any cluster; its expression remained remarkably unaffected by ERK inhibition up to 8 hpa and was slightly decreased at 12 hpa, although not meeting our significance threshold (Fig. 5B).

Fos, a bZIP TF and IEG, is a known phosphorylation target of ERK in mammals^21^ and like Jun, may increase its transcription through autoregulation.^51,52^ Furthermore, bZIP TFs, such as Fos, Jun and CREB were previously implicated in regulating Wnt gene transcription in *H. vulgaris,* and *fos jun*, and *creb* transcripts are upregulated as part of the generic injury response.^27,36^ We therefore asked whether ERK inhibition affects bZIP TF transcript levels. We observed no changes in *jun* and *fos* transcript levels up to 8 hpa in response to ERK inhibition, with only a slight reduction at 12 hpa that did not meet our significance threshold, and *creb* transcript levels remained unchanged throughout the 12-hour treatment (Fig. 5C). These results indicate that ERK inhibition does not affect their rapid transcriptional upregulation following injury, although this does not exclude the possibility that ERK regulates these factors post-transcriptionally through phosphorylation. On the other hand, we did observe transcriptional downregulation of the ETS family TF *elk-1* (G004172) up to 12 hpa (Fig. 5C), revealing a previously less appreciated role for this TF in the early injury response in *H. vulgaris*. Several other ETS family TFs were unaffected by ERK inhibition (Supp. Fig. 10B).

### ERK inhibition reduces the spread of the injury domain from the wound edge

Injury is known to trigger the propagation of ERK signaling waves across tissue, such as in mouse epithelial injury, planarian whole body regeneration, or zebrafish scale regeneration.^12,18,53^ These ERK signaling waves are thought to transmit the injury signal away from the wound site and coordinate a regenerative response. Therefore, we next asked if dampening ERK signaling with U0126 during *Hydra* head regeneration reduces the spread of the injury signal, resulting in a smaller expression domain of injury-induced transcription.

To assess how ERK inhibition affects the spread of injury-induced transcripts from the wound edge, we quantified the area of injury-responsive gene expression in whole-mount regenerating *Hydra* heads using Fluorescence *in Situ* Hybridization (**FISH**), normalizing measurements to the width of the body column. At 12 hpa, *wnt3, wnt9/10c, wntless*, *jun,* and *fos* were expressed from the wound edge into the body column tissue (Fig. 6A, Supp. Fig. 12, and Supp. Fig. 13). In response to ERK inhibition, the expression domain area was significantly retracted as compared to controls for all five genes at 12 hpa (Figs. 6A-C, Fig. 6E, Supp. Fig. 12, and Supp. Fig. 13).

**Fig. 6.**
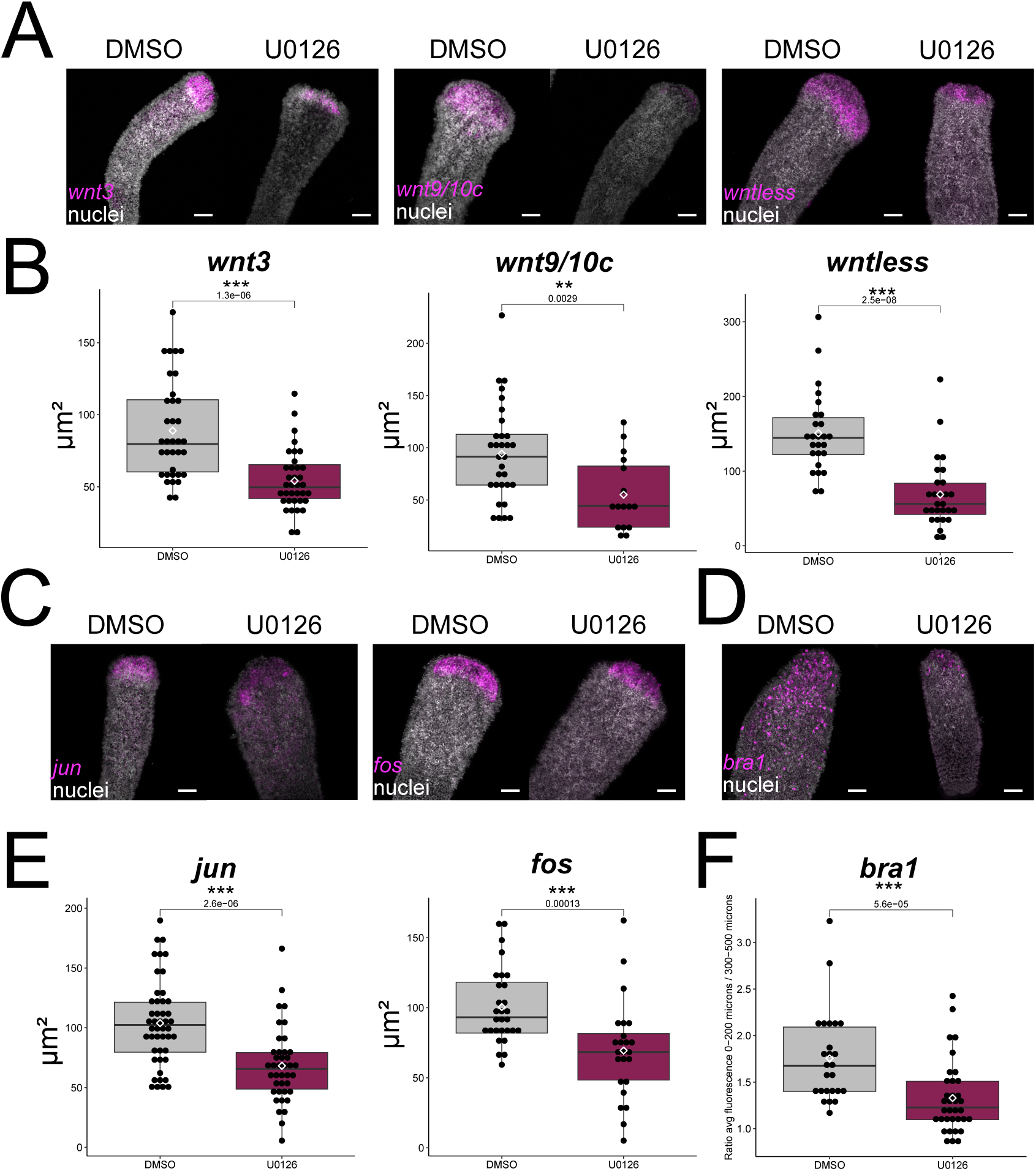
ERK inhibition reduces the area of the injury domain. (A) Representative images of regenerating head tips treated with DMSO or U0126 for the first 12 hours of regeneration, collected for fluorescence *in situ* (FISH) (magenta) and Hoechst nuclear staining (white). Scale = 100µm. Sample sizes are as follows: *wnt3* (DMSO n=35, U0126 n=36), *wnt9/10c* (DMSO n=31, U0126 n=16), and *wntless* (DMSO n=26, U0126 n=28). (B) Quantification of expression domain area for *wnt3, wnt9/10c,* and *wntless* comparing DMSO controls (gray) and U0126-treated animals (magenta). Expression area was normalized by body column width. Each point represents one animal, and the diamond indicates the mean. Statistical comparisons were performed using the Wilcoxon rank-sum test. (C,D) Representative images of regenerating head tips treated with DMSO or U0126 for the first 12 hours of regeneration, collected for FISH (magenta) and Hoechst nuclear staining (white). Scale bar = 100µm. Sample sizes are as follows: *jun* (DMSO n=47, U0126 n=40), *fos* (DMSO n=29, U0126 n=24), and *bra1* (DMSO n=25, U0126 n=35). (E) Quantification of expression domain area for *jun* and *fos* comparing DMSO controls (gray) and U0126-treated animals (burgundy). Expression area was normalized by body column width. Each point represents one animal, and the diamond indicates the mean. Statistical comparisons were performed using the Wilcoxon rank-sum test. (F) Quantification of *bra1* expression from the wound edge comparing DMSO controls (gray) and U0126-treated animals (burgundy). Each point represents one animal, and the diamond indicates the mean. Statistical comparisons were performed using the Wilcoxon rank-sum test.

While *wntless* expression was significantly decreased at 12 hpa in our RNA-seq data, *wnt3*, *wnt9/10c, jun*, and *fos* were not (Figs. 5B and C). However, all four showed a downward trend at 12 hpa, and importantly, our spatial analysis measures expression area rather than transcript intensity. Thus, even when absolute expression levels are only modestly reduced, the spatial domain of expression can still contract.

Finally, consistent with our RNA-seq data (Fig. 5B), there is a retraction of the *bra1* expression domain (Figs. 6D and F). Notably, *bra1* is not expressed uniformly across the injury domain and instead shows strong expression in scattered cells, consistent with its established expression in gland cells, observed in previous work.^34,48^ In uninjured *Hydra*, *bra1* is comparatively uniformly expressed across the hypostome, in the ectodermal and endodermal epithelial cells, as well as in oral gland cells, a pattern that is consistent with single-cell transcriptomic profiles and prior in situ observations (Supp. Fig. 14).^34,48,54^ For reasons that are not yet clear, injury-induced Wnt signaling appears to transiently upregulate *bra1* expression in gland cells as compared to surrounding epithelial cells at the early stages of regeneration. Regardless, our data show that the area of Wnt-responsive gene expression is reduced following ERK inhibition, consistent with the reduced area of Wnt ligand expression. Overall, we find evidence that ERK signaling is required to establish the full size of the transcriptional injury domain in regenerating *H. vulgaris* heads, suggesting a role for ERK in propagating the injury signal through the tissue.

### Distinct ERK signaling dynamics in H. oligactis and H. vulgaris are associated with divergent regenerative phenotypes

*Hydra oligactis* differs from *H. vulgaris* in regenerative ability: it regenerates its head more slowly, and has lower foot regeneration success.^40,41^ We recently showed that *H. oligactis* has a strongly reduced injury-induced Wnt transcription response, contributing to its foot regeneration deficiency.^41^ Wnt expression in the regenerating head also rises more slowly, reaching strong levels only by 24 hpa, delaying oral organizer formation compared to *H. vulgaris*, where robust Wnt activation and organizer formation occur between 8 and 12 hpa.^34,36,55,56^ Given our finding that ERK signaling supports robust injury-induced Wnt activation in *H. vulgaris*, we next examined ERK signaling dynamics and function in *H. oligactis* to determine whether differences in this pathway underlie their divergent regenerative abilities.

We first tested injury-induced p-ERK signaling dynamics in *H. oligactis* using immunofluorescence and observed high p-ERK signal at the wound edge at 0.5 hpa, similar to *H. vulgaris* (Fig. 7A). However, while p-ERK signal at the wound edge gradually decreased over the first 12 hpa in *H. vulgaris*, it rapidly dropped to a near-homeostatic level by 3 hpa in *H. oligactis* and then remained at this slightly elevated level through 48 hpa (Fig. 7A-C, Supp. Fig. 15, and Supp. Fig. 16). Similarly, the distal wound in *H. oligactis* showed a rapid drop in p-ERK by 3 hpa (Supp. Fig. 2B). In summary, our experiments show that p-ERK levels in *H. oligactis* fall to near-homeostatic levels more rapidly as compared to *H. vulgaris*, which correlates with the lack of a strong injury-induced upregulation of Wnt signaling in *H. oligactis*.^41^

**Fig. 7.**
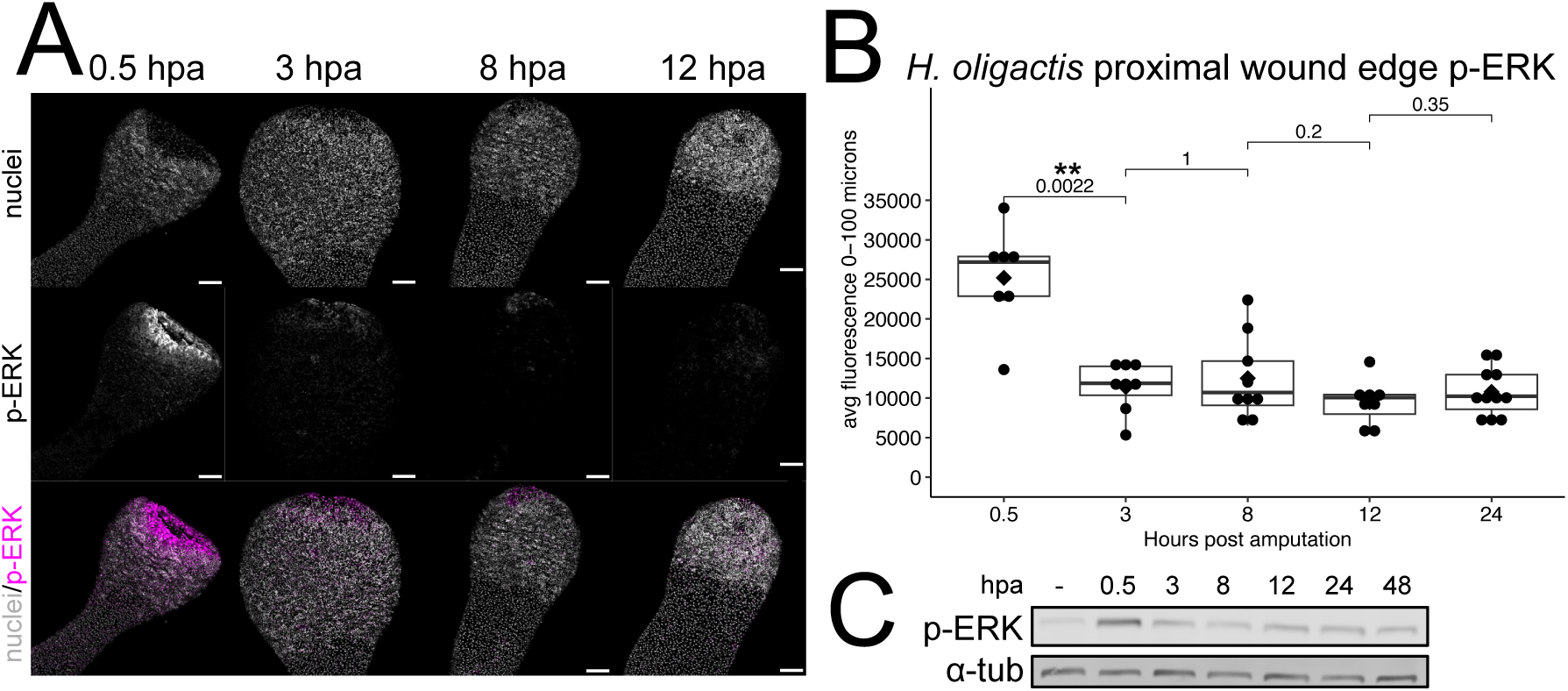
*Hydra oligactis* displays a brief injury-induced phosphorylation of ERK. (A) Immunofluorescence of regenerating *H. oligactis* heads using an antibody against p-ERK together with Hoechst to label nuclei shows p-ERK signal at the wound edge that is strong at 0.5 hpa and then rapidly dissipates. Scale bar =100µm. (0.5 hpa n=7, 3 hpa n=8, 8 hpa n=9, 12 hpa n=8, 24 hpa n=11) (B) Quantification of the experiment in panel A. Each point represents one animal. Intensity was measured using plot profile. Statistical comparisons were performed using the Wilcoxon rank-sum test. (C) Western Blot showing p-ERK and α-tubulin (loading control) protein levels during *H. oligactis* head regeneration. The first lane shows the basal level of p-ERK in whole animals.

We next tested the effect of inhibiting ERK signaling on head regeneration in *H. oligactis.* We confirmed that homeostatic *H. oligactis* is tolerant to long-term U0126 treatment at 25µM (Supp. Fig. 17) and that injury-induced p-ERK is dampened by U0126 treatment (Fig. 8A-B and Supp. Fig. 18A). We next compared the effects of continuous U0126 treatment on head regeneration in *H. vulgaris* and *H. oligactis*. Whereas sustained ERK inhibition caused a nearly complete block of head regeneration in *H. vulgaris* up to 96 hpa, almost half of the treated *H. oligactis* polyps regenerated heads by this time (Fig. 8C). Between 72 hpa and 96 hpa, U0126 treated animals started to exhibit cell sloughing and death in both species, consistent with disruption of essential homeostatic processes, therefore experiments were stopped at 96hpa (Supp. Fig. 18B). The increased ability of *H. oligactis* to regenerate its head under continuous ERK inhibition suggests a weaker dependence on ERK signaling for activating Wnt pathways. However, transient ERK inhibition during the first 12 hpa delays head regeneration as compared to DMSO controls in *H. oligactis*, though the effect is less pronounced than in *H. vulgaris* (Fig. 8D and Supp. Fig. 18C). Together, these findings highlight clear differences in ERK signaling function between the two species that could influence their different regenerative phenotypes.

**Fig 8.**
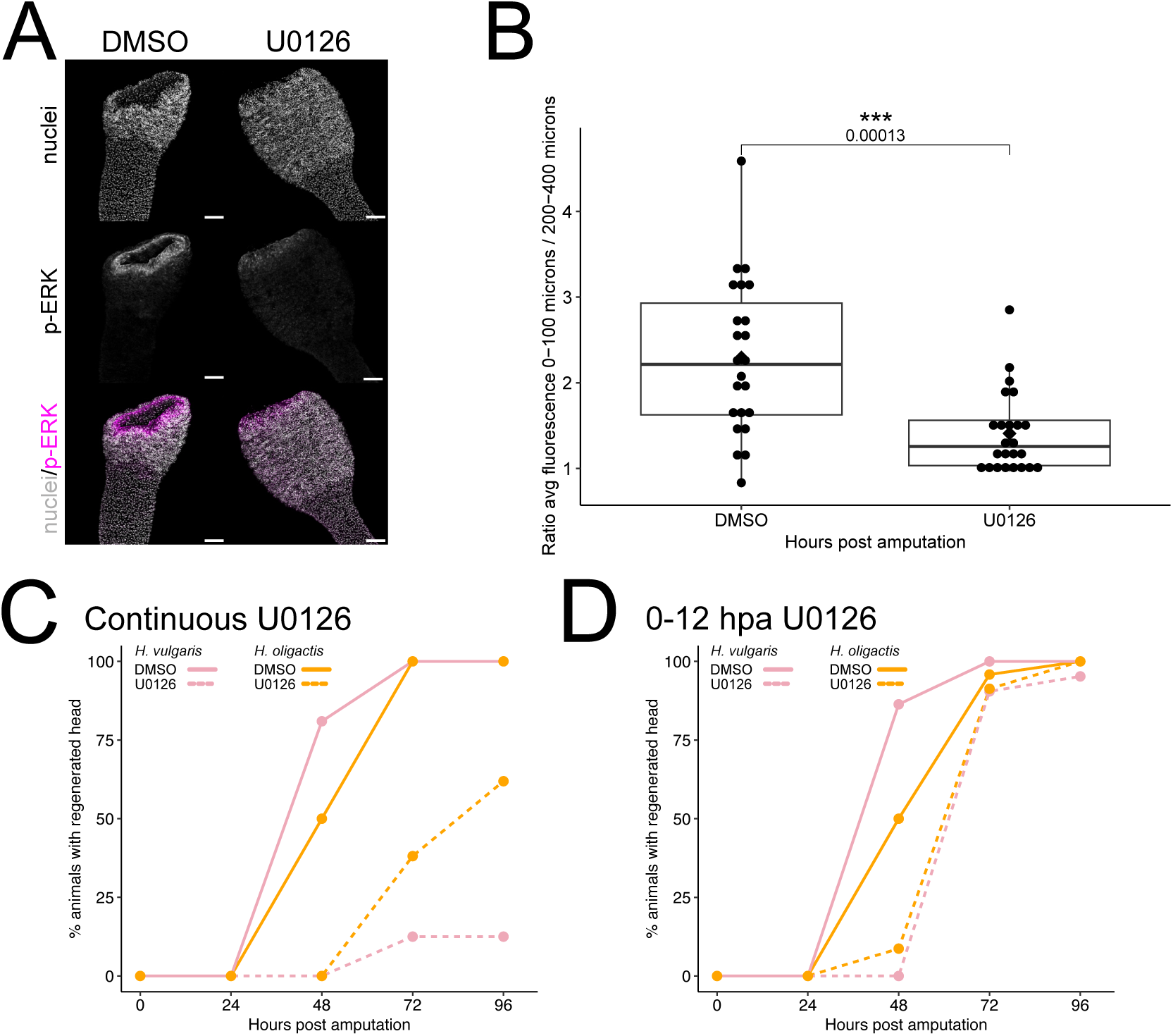
Head regeneration in *Hydra oligactis* is less sensitive to ERK signaling inhibition. (A) Immunofluorescence of regenerating *H. oligactis* heads at 0.5, 3, 8 and 12 hpa, treated with DMSO or 25µM U0126, using an antibody against p-ERK together with Hoechst to label nuclei. Scale bar = 100µm. (B) Quantification of *H. oligactis* p-ERK enrichment at the wound edge from the experiment in panel A. (C) Comparison of head regeneration rates between *H. vulgaris* and *H. oligactis* treated continuously with DMSO vehicle control or 25µM U0126. (D) Comparison of head regeneration rates between *H. vulgaris* and *H. oligactis* treated with DMSO or 25µM U0126 during the first 12 hours of regeneration. For C and D, solid lines represent DMSO treatment, and dotted lines represent U0126 treatment. Orange lines represent *H. oligactis* and pink lines represent *H. vulgaris*.

## DISCUSSION

The activation of ERK signaling following injury is deeply conserved across animals,^6,16,17,27,57–60^ yet how this signal is translated into a regenerative response, or fails to be, remains an open question. MAPK pathways commonly activate IEG TFs, including bZIP and EGR family members,^21^ and in some systems, such as the *Drosophila* wing disc and acoel regeneration, these TFs directly induce transcription of Wnt signaling genes that are crucial for patterning regenerated tissue.^19,23,61^ However, the structure of the GRNs driving regeneration and the mechanisms linking injury to pattern formation remain incompletely understood. In this study, extending previous findings that injury-induced ERK signaling promotes *H. vulgaris* head regeneration,^27^ we refine the temporal framework of ERK activation and show that its activity continues beyond the generic injury phase into the pre-patterning phase (Fig. 9A). ERK is required for the early activation of several Wnt pathway components, although *wnt3* is initially resistant to perturbation, indicating that distinct Wnt ligands are regulated through different mechanisms. ERK signaling is also necessary for expanding the transcriptional injury domain, and its dynamics differ markedly in *H. oligactis*, where p-ERK attenuates rapidly, and regeneration is correspondingly less sensitive to inhibition. Together, these findings clarify ERK’s role in early patterning and suggest that evolutionary variation in the persistence of ERK signaling, contributes to differences in regenerative capacity.

**Fig. 9.**
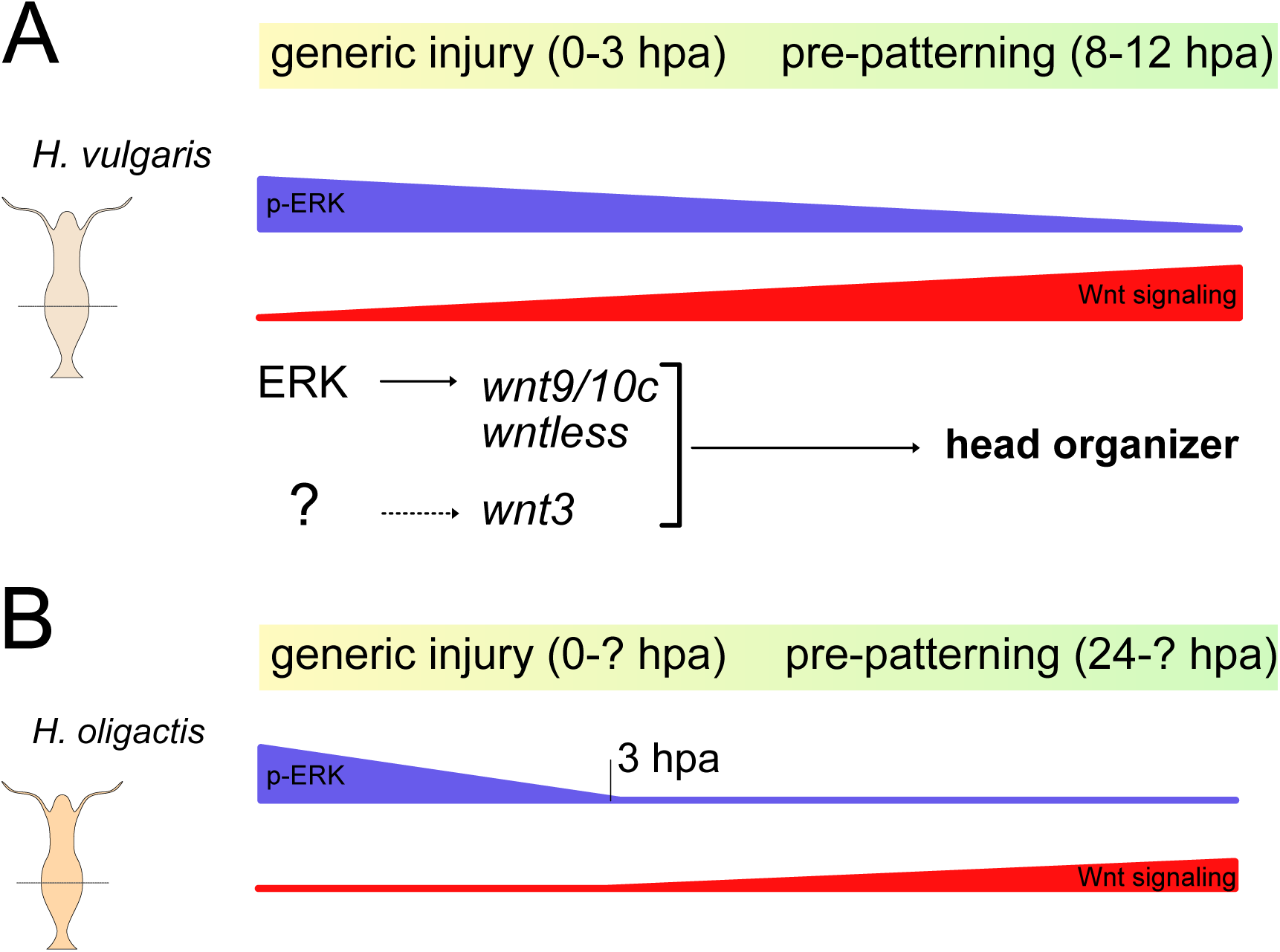
Comparative model illustrating divergent ERK signaling roles in head regeneration in *H. vulgaris* and *H. oligactis*. (A) Model of ERK and Wnt signaling dynamics during head regeneration in *H. vulgaris.* P-ERK levels decline gradually over the first 12 hours after injury, while the transcription of Wnt signaling components and the associated head organizer activity increase over this same period. Transcription of *wnt9/10c* and *wntless* requires ERK signaling, while *wnt3* transcription may be activated by a distinct injury-induced signal. (B) Model of ERK signaling and Wnt signaling dynamics during head regeneration in *H. oligactis.* p-ERK levels drop sharply before 3 hpa, and the onset of Wnt signaling upregulation is comparatively delayed.^41^

In our previous work, we showed that the generic response to injury in *H. vulgaris* includes the transcriptional activation of the bZIP TFs *jun*, *fos*, and *creb*.^36^ ATAC-seq analysis further revealed that CRE motifs, which are bound by bZIP TFs such as Jun, Fos, and CREB, are present within several Wnt pathway genes, including *wnt3*, *wnt9/10c*, and *wntless*, and that these sites become more accessible after injury.^36^ These observations support the hypothesis that injury-induced bZIP TFs contribute to Wnt gene activation. Given the conserved role of ERK signaling in activating bZIP TFs,^21,62,63^ it is plausible that the downregulation of Wnt signaling genes following ERK inhibition operates, at least in part, through bZIP TF-dependent mechanisms. However, we did not observe strong disruption of *jun*, *fos*, or *creb* transcription in response to ERK inhibition, which is somewhat unexpected because these TFs often participate in feedforward loops that enhance their own transcription.^51,52^ This result does not rule out a post-translational requirement for ERK in activating bZIP TFs during *H. vulgaris* regeneration, which is supported by a previously conducted proteomics study showing enrichment for the ERK phosphorylation motif between 3-12 hpa, but will require additional proteomic and biochemistry approaches to more clearly unravel.^64^ By contrast, ERK inhibition clearly reduced expression of the ETS TF *elk1*, a known ERK target in other systems.^21,65,66^ Our earlier work suggested that ETS TFs contribute to injury-induced upregulation of *wntless*,^36^ and the current data places ERK upstream of this activity, as *elk1* and *wntless* are both downregulated when ERK signaling is inhibited. Together, these results support a model in which reduced ERK signaling impacts the expression of Wnt signaling genes and at least some of their likely upstream TFs.

We found that ERK inhibition did not immediately affect *wnt3* transcript levels; they remained stable through 8 hpa and showed only a slight decline at 12 hpa. This delayed response suggests that *wnt3* is regulated differently from other Wnt pathway components and may not rely directly on ERK signaling (Fig. 9A). Its later downregulation may reflect indirect consequences of earlier disruptions. Prior work from the Holstein lab reported only mild transcriptional effects of ERK inhibition on *wnt3* and *wnt9/10C* using qPCR,^27^ whereas our RNA-seq data provide higher sensitivity and reveal gene-specific differences more clearly. Our results suggest that *wnt3* activation alone is insufficient for timely head regeneration. However, the activity of Wnt3 protein could be affected by the reduced expression of *wntless*, which is required for Wnt ligand secretion and significantly downregulated in our data set.^67^ We further propose that the regulation of *wnt3* differs from that of *wnt9/10C* and *wnt7*. Supporting this idea, previous qPCR studies showed that inhibitors of p38 and JNK MAPK pathways upregulate *wnt3* while downregulating *wnt9/10c*, reinforcing that these Wnt ligands respond differently to injury-associated signals.^27^ How *wnt3* expression is activated following injury therefore remains an open question, and its distinct regulatory behavior underscores the complexity of the Wnt module during early regeneration.

Transient inhibition of ERK signaling leads to a stable block of planarian whole-body and zebrafish tail fin regeneration that can be reversed by inflicting a new injury.^17^ This contrasts sharply with *H. vulgaris*, where transient ERK inhibition spanning the entire generic injury phase causes a delay but not an arrest of head regeneration. In the planarian *Schmidtea mediterranea*, an initial burst of generic injury-induced proliferation is ERK independent, but ERK activity is subsequently required to drive the proliferation necessary for blastema formation in regenerative wounds.^17^ Likewise, whole-body regeneration in acoels is inhibited by transient ERK inhibition, coincident with a significant reduction in proliferation.^19^ A similar reliance on ERK-driven proliferation is seen in the cockroach, where ERK activates casein kinase 2 (CK-2) to promote blastema formation during appendage regeneration.^14^ *Hydra*, however, regenerates in a different proliferative context. Epithelial cells in the body column are already cycling prior to injury, providing a preexisting pool of proliferative cells that allows regeneration to move directly from the generic injury response into the initiation of patterning.^29^ This distinction effectively decouples ERK’s role in proliferation from its role in triggering patterning programs in *Hydra* and allows us to focus specifically on how ERK regulates injury-induced transcriptional events, including the activation of Wnt pathway components. Consistent with this interpretation, our differential gene expression analysis revealed no clear downregulation of proliferation-associated genes in regenerating or homeostatic animals treated with U0126, suggesting that ERK plays little or no direct role in maintaining proliferation in *Hydra*.

Injury-induced ERK signaling is known to propagate outward from the wound site, helping communicate the wounded state across the tissue.^12,18,19,53^ In several systems, this propagation occurs as ERK signaling waves that transmit information from proximal to distal regions, allowing both nearby and distant tissues to participate in coordinating regeneration across large distances.^12,18^ In *H. vulgaris*, we likewise observed elevated p-ERK levels in tissue both proximal and distal to the wound. To test how this propagation affects the transcriptional response to injury, we examined how ERK inhibition affects the spatial expression patterns of genes activated during the generic injury response. The expression domains of *wnt3*, *wnt9/10c*, *wntless*, *jun*, *fos*, and *brachyury1* were all retracted toward the wound edge when ERK signaling was inhibited, indicating that ERK is required for the transcriptional response to extend deeper into the tissue. This suggests that ERK plays a key role in relaying injury signals across the regenerating body column. The mechanisms underlying this propagation likely involve upstream signals such as ROS and Ca²⁺, which are known to modulate MAPK signaling strength in *H. vulgaris*.^27^

The rapid recovery of p-ERK following U0126 washout we observe in *H. vulgaris* suggests that upstream injury signals persist during inhibition and are poised to re-activate ERK as soon as the inhibitor is removed. Such signals could involve ROS or Ca²⁺, although this remains to be directly tested. The differing regeneration outcomes after 3-hour versus 12-hour transient inhibition offer additional clues. After a 3-hour treatment, regeneration proceeds with essentially normal timing, raising the possibility that upstream injury cues accumulate while ERK is blocked and produce a strong rebound once inhibition is lifted. In contrast, 12 hours of inhibition introduces a roughly 24-hour delay in regeneration, consistent with a scenario in which prolonged ERK suppression slows or prevents the timely amplification of Wnt signaling. This delayed recovery resembles the naturally slower regeneration dynamics of *H. oligactis*, suggesting that reduced or transient ERK activity pushes *H. vulgaris* into a regenerative mode more characteristic of *H. oligactis*.

The striking divergence in ERK signal duration between *H. vulgaris* and *H. oligactis* offers a compelling explanation for their contrasting regenerative capacities. Although both species mount a strong p-ERK response immediately after injury, *H. vulgaris* sustains high signaling at the wound site, whereas p-ERK levels in *H. oligactis* drop sharply and return close to baseline by 3 hpa (Fig. 9). This difference aligns with a broader pattern seen across regenerative animals, in which regenerative competence correlates more strongly with the duration of ERK signaling than with its initial activation. Regeneration-competent muscle cells in salamanders show sustained ERK activation, allowing re-entry of the cell cycle; conversely, less regenerative mammalian cells only transiently activate p-ERK.^16^ A similar dichotomy exists between *Acomys* and *Mus*; the regenerative spiny mouse sustains ERK activity, whereas the non-regenerative laboratory mouse exhibits only brief activation, resulting in scarring.^6^ Even in planaria, where pERK is induced by any injury, regeneration but not wound healing requires ongoing ERK signaling.^17^

Our results suggest that the evolution of regenerative ability in *Hydra* is driven, at least in part, by modulation of ERK signaling, aligning with results in distantly related animals. The sustained ERK phase in *H. vulgaris* appears necessary to drive the rapid, injury-induced Wnt transcriptional burst that underlies its robust regenerative ability, whereas the slower timeline of head regeneration in *H. oligactis* correlates with its inability to maintain high-level ERK signaling beyond the initial injury-induced burst (Fig. 9B). This interpretation is supported by functional perturbation experiments: unlike *H. vulgaris*, which largely fails to regenerate under continuous U0126 treatment, a substantial fraction of *H. oligactis* eventually regenerates despite the inhibition. This resilience implies that *H. oligactis* relies on an alternative, slower mechanism for establishing the head organizer, either independent of high ERK activity or requiring only a minimal ERK threshold.

Viewed this way, we propose that the sustained ERK-Wnt module in *H. vulgaris* functions as a regeneration “accelerator.” Transient ERK inhibition removes this accelerator, forcing *H. vulgaris* into a slower, *H. oligactis*-like regenerative mode. As a consequence, regeneration rates in the two species converge under transient inhibition. Consistent with this framework, our previous work showed that transient inhibition of Wnt signaling slows *H. vulgaris* regeneration to match that of *H. oligactis*, while Wnt agonist treatment accelerates regeneration in *H. oligactis*,^41^ further supporting the idea that regeneration speed is tuned by the strength and persistence of early injury-induced patterning signals.^6,17,68,69^

## CONCLUSION

Together, our results support the emerging view that ERK is not merely an injury trigger but a sustained, spatially propagating, GRN-shaping signal whose duration, tissue spread, and regulatory specificity collectively influence regenerative outcomes. The contrasting ERK dynamics of *H. vulgaris* and *H. oligactis* illustrate how the persistence of early injury signaling can be evolutionarily tuned in ways that modulate regenerative capacity. In *Hydra*, this clarity is enhanced by the ability to dissect regulatory interactions among injury signals, ERK, and Wnt signaling without the confounding influence of blastema-dependent proliferation, allowing us to identify conserved nodes that anchor regeneration GRNs across species. Such nodes offer traction for comparing regeneration mechanisms even at large evolutionary distances where tissue contexts and injury responses differ markedly.^9^ Looking forward, systematic mapping of regeneration GRNs with high resolution comparable to that achieved in developmental systems will be essential for constructing a unified cross-species framework that explains how regenerative abilities evolve, are lost, and are regained.^7,70–72^ Our study provides new building blocks toward this broader comparative understanding of how regeneration is orchestrated and has evolved across the animal kingdom.

## METHODS

### Code and Data Availability

All analysis code is available in the git repository (https://github.com/cejuliano/ERK-Signaling-head-regeneration-in-Hydra).

Raw RNA-seq FASTQ files have been deposited under accession PRJNA1373231. *Hydra* AEP gene models are available through the *Hydra* AEP Genome Project Portal: (https://research.nhgri.nih.gov/HydraAEP/).^73^

### Hydra vulgaris and Hydra oligactis culture

The strains used in this study were *Hydra vulgaris* AEP^36^ and *Hydra oligactis* Innsbruck 12.^41^ The *Hydra* strains used in this study were cultured in *Hydra* Medium (HM) (0.38 mM CaCl_2_, 0.32 mM MgSO_4_ X 7H_2_O, 0.5 mM NaHCO_3_, 0.08 mM K2CO_3_) at 18°C under a 12 hr light-dark cycle. *Hydra* were fed freshly hatched, premium grade *Artemia* nauplii (https://www.brineshrimpdirect.com) three times per week and were transferred into fresh HM 4-8 hours after feeding. Animals were starved for 24 hours prior to experimentation.

### ERK inhibitors

U0126 (Cell Signaling, #9903S) and BVD-523 (MedChemExpress, #HY-15816) were resuspended in DMSO at 25 mM and 7.5 mM, respectively, then aliquotted and stored at -70°C. Aliquots were thawed at room temperature and leftovers were discarded after use. For all pharmacological experiments, DMSO (vehicle control) was diluted 1:1000 in HM. U0126 was used at 25 µM in HM and BVD-523 was used at 7.5 µM in HM unless otherwise indicated.

Diluted inhibitors were mixed thoroughly by vigorous shaking in a conical tube. Following addition of inhibitors to HM, animals were incubated at 18°C.

### Head Regeneration Assays

Whole, bud-free *Hydra* were placed in 6-well plates with 3mL HM containing DMSO (1:1000), 25 µM U0126, or 7.5 µM BVD-523 for one hour prior to transverse bisection at the midpoint of their oral-aboral axis with a scalpel (Word Precision Instruments, #500239). For the regeneration assay in Figure 2, animals remained in the drug treatment until the indicated time, after which all groups except the 0-12 hpa treatment were washed three times in HM containing DMSO (1:1000). At 12 hpa, all animals were washed three times in drug-free HM. For the regeneration assay in Figure 8C, animals remained continuously in drug, which was refreshed every 24 hours. For the regeneration assay in Figure 8D, animals regenerated in drug until 12 hpa and were then washed three times in drug-free HM.

At 24 hours post bisection, each regenerating head was transferred to an individual well of a 12-well plate containing 1 mL HM. Animals were imaged using a Leica 582 DFC300 G camera and LAS v4.7 software, and tentacle buds were counted. Tentacle counts were repeated every 24 hours.

### Western Blot Assays

Western blots were performed following a previously published protocol with minor modifications.^27^ For untreated whole-animal samples, seven bud-free *H. vulgaris* were transferred to a 1.5mL tube with minimal media carryover. For drug-treated samples, 15 bud-free *H. vulgaris* were incubated in DMSO or 25 µM U0126 for one hour prior to transverse bisection at the midpoint of their oral-aboral axis. Regeneration proceeded for the indicated time in the drug treatment before tissues were collected or washed in HM. Regenerating heads were transferred to 1.5mL tubes with minimal excess medium. Experiments including *H. oligactis* followed the same protocol, except that five whole animals were collected for baseline p-ERK levels and 10 regenerating heads were collected for each regeneration timepoint.

PhosSTOP (Roche, #4906845001) and cOmplete (Roche, #11697498001) were added to the lysis buffer (4% w/v SDS; 0.5% w/v sodium deoxycholate; 20 mM Tris, pH 7.5). 50µL of lysis buffer was added to each sample, followed by heating at 95°C for 10 minutes with intermittent vortexing. Lysates were transferred to clean 1.5mL tubes and cooled on ice.

Protein concentration was measured using the Pierce BCA Assay Kit (Thermo Scientific, #23225) with the enhanced protocol (1mL working reagent; 50 uL of 10x diluted lysate).10µg of protein was mixed with 4X sample loading buffer (0.2 M Tris-HCl, 0.4 M DTT, 8% (w/v) SDS, 6 mM Bromophenol blue, 4.3 M glycerol) and heated at 95°C for 10 minutes. After cooling to room temperature, samples were separated on Bolt Bis-Tris Plus 4-12% Mini Protein Gels (Invitrogen, #NW04122BOX) with Bolt MES SDS buffer (Invitrogen, #B0002) at 200 Volts for 25 minutes.

Proteins were then transferred to a 0.2 µm nitrocellulose membrane (Invitrogen, #LC2000) at 30mA for 16 hours at 4°C in chilled buffer (10% v/v 10x running buffer; 20% v/v methanol) with an ice pack.

Membranes were washed 3X in TBST and blocked for 2 hours in blocking buffer (5% v/w BSA, TBST, 5 mM EDTA, pH 8.0, 1 mM EGTA) at room temperature. After three additional TBST washes, membranes were incubated overnight at 4°C with primary antibodies (1:1000 mouse anti-tubulin: Sigma-Aldrich, #T6199; 1:1000 rabbit anti-phospho-ERK: Cell Signaling, #4370) diluted in the blocking buffer. After three TBST washes, membranes were incubated with secondary antibodies (1:5000 568-goat anti-mouse, Thermo Fisher, #A-11004; 1:2000 488-goat anti-rabbit, Thermo Fisher #A-11008), in blocking buffer for 1.5 hours at room temperature. Final washes (3× TBST) preceded imaging on a Sapphire Azure Scanner.

### RNA-seq sample collection

RNA-seq sample preparation followed a previously published protocol.^36^ In brief, for the regeneration dataset, 30 bud-free *Hydra vulgaris* polyps that had been fed three times weekly were starved for 24 hours, then treated with DMSO or 25 µM U0126 for 1 hour. Animals were then transversely bisected at the midpoint of their oral-aboral axis in batches of five to minimize variability in amputation timing. Regeneration proceeded for 0, 3, 8, or 12 hours in the drug treatment. Regenerating head tissue (approximately one-third of the regenerating animal’s length) was isolated and frozen in Trizol (Invitrogen, #15596018) at -70°C. Six batches of five regenerating heads were pooled to generate one biological replicate. For the whole, uninjured *Hydra* dataset, 10 bud-free *Hydra vulgaris* polyps that had been fed three times weekly were starved for 24 hours, then treated with DMSO or 25 µM U0126 for 12 hours. For each treatment and timepoint, three biological replicates (totaling 30 regenerating animals or 10 whole animals per replicate) were prepared, following the standard practice of using three replicates for high-throughput sequencing. Samples were submitted to Novogene for RNA quality assessment, library preparation, and sequencing. RNA quantity and integrity was assessed using 1% agarose gel electrophoresis, NanoDrop spectrophotometry, and Agilent 2100 gel electrophoresis. mRNA libraries were prepared using polyA enrichment and sequencing was performed on a NovaSeq 6000 platform using 2 x 150 bp paired-end reads.

### RNA-seq data processing

RNA-seq data were processed using a previously published pipeline,^36^ with the exception that reads were mapped to the updated *Hydra vulgaris* AEP gene models (https://research.nhgri.nih.gov/HydraAEP/). Sequencing adapters and low-quality base calls were removed from the raw sequencing data using Trimmomatic v 0.36. Filtered reads were mapped to the *Hydra vulgaris* AEP gene models, and read counts per gene were calculated using RSEM.^74^ Full processing details and scripts are available in the GitHub repository (https://github.com/cejuliano/ERK-Signaling-head-regeneration-in-Hydra).

### RNA-seq data analysis

The RNA-seq libraries were obtained in two different batches, therefore we first corrected for batch effects using the ComBat_seq R-package^75^ to create a merged transcript-count matrix. Batch-corrected counts were then normalized, and differential gene expression analysis was performed using edgeR2.^76^ Differentially expressed genes (DEGs) were identified with glmTreat, using a log2-fold change threshold of 1.1 and a false discovery rate (FDR) of 0.05. DEG tables are provided in Table 1. Time-course expression plots were generated using custom scripts provided in the GitHub repository (https://github.com/cejuliano/ERK-Signaling-head-regeneration-in-Hydra).

Temporal patterns of differential gene expression during head regeneration were analyzed using maSigPro,^49^ an R package that identifies genes with significant expression differences between conditions across time using a regression-based framework. Parameters followed recommended settings for normalized counts per million (CPM) values (theta = 10, Q = 0.05, MT.adjust = “BH”, alfa = 0.05) and clusters were filtered with a conservative R-square threshold of 0.8. This analysis grouped 306 genes into nine clusters with distinct temporal differential gene expression patterns between U0126 and control conditions.

We performed Gene Ontology (GO) enrichment analysis to test for enrichment of GO terms in the Biological Processes domain using the topGO R package (Alexa A, Rahnenführer J (2025). *topGO: Enrichment Analysis for Gene Ontology*. doi:10.18129/B9.bioc.topGO, R package version 2.62.0, https://bioconductor.org/packages/topGO.), with enrichment p-values computed by Fisher’s exact test and adjusted for multiple testing using the Benjamini–Hochberg method (FDR < 0.05). Redundant GO enriched terms were collapsed using MonaGO by Resnik GO term similarity.^77^ GO-term dot plots were generated using custom scripts provided in the GitHub repository. Gene Set Enrichment Analysis (GSEA 4.4.0) was used to assess enrichment of Kyoto Encyclopedia of Genes and Genomes (KEGG) and the Reactome Pathway Database.

### Whole-mount immunofluorescence for p-ERK detection

The protocol was adapted from a previously published method with minor modifications.^27^ Samples were incubated in 2% urethane HM for one minute in 1.5mL tubes. Samples were then washed briefly with freshly diluted 4% formaldehyde in PBST, then the samples were placed into fresh fixative, and incubated overnight at 4 °C with gentle rocking.

The following day, samples were washed three times in TBST and permeabilized in 0.1% v/v Triton-X 100 in TBS for 15 minutes, followed by three TBST washes. Tris-based antigen retrieval (Inoue) was performed by incubating samples with 150 mM Tris-HCl at pH 9.0 for 5 minutes at room temperature and then heating at 70°C for 15 minutes. After three additional TBST washes, samples were incubated in blocking buffer for 2 hours at room temperature (5% v/w BSA, TBST, 5mM EDTA, pH 8.0, 1 mM EGTA). Samples were incubated in primary anti-p-ERK antibody (Cell Signaling, #4370) at a 1:100 dilution in blocking buffer (1% v/w BSA, TBST, 5mM EDTA, pH 8.0, 1 mM EGTA) and incubated at 4°C overnight.

The next day, samples were washed three times with TBST and incubated in secondary antibody (Thermo Fischer, Goat anti-Rabbit IgG Cross-Adsorbed Alexa Flour 568, #A-11004) at 1:500 in blocking buffer (1% v/w BSA, TBST, 5mM EDTA, pH 8.0, 1 mM EGTA) in the dark for 2.5 hours at room temperature. Samples were then washed three times with TBST, with the second wash containing 1:1000 Hoechst (Thermo Fischer, #H3570). Finally, samples were passed through a glycerol series before mounting in 80% glycerol with a spacer (EMS #703279S) for imaging.

### Whole-mount immunofluorescence for phospho-histone-3 detection

Animals were pre-treated for one hour with 1:1000 DMSO, 25 μM U0126, or 7.5 μM BVD-523 before bisection. Regenerating head samples were directly incubated in 2% urethane HM for one minute in 1.5mL tubes. Samples were fixed at room temperature in freshly diluted 4% formaldehyde with gentle rocking for one hour. Following fixation, samples were washed three times in PBST and solubilized in 0.1% v/v Triton-X 100 in PBS for 15 minutes. After three additional washes in PBST, samples were incubated in blocking buffer for 2 hours at room temperature (1% BSA w/v, 10% goat serum v/v; 0.1% Triton X-100 in PBS). Primary antibody incubation was performed overnight at 4°C anti-pH3 antibody (Cell Signaling, #4370) diluted 1:200 in blocking buffer (1% BSA w/v, 10% goat serum v/v; 0.1% Triton X-100 in PBS).

The next day, samples were washed three times with PBST and incubated in secondary antibody (Thermo Fischer, Goat anti-Rabbit IgG Cross-Adsorbed Alexa Flour 568, #A-11004) diluted 1:500 in blocking buffer (1% BSA w/v, 10% goat serum v/v; 0.1% Triton X-100 in PBS) in the dark for 2 hours at room temperature. Samples were washed three additional times with PBST, with the second wash containing 1:1000 Hoechst (Thermo Fischer, #H3570). Samples were passed through a glycerol series before mounting in 80% glycerol with a spacer (EMS #703279S) for imaging.

### Probe Generation for Fluorescent in Situ Hybridization

Labeled RNA probes were generated from oligo-dT15 primed cDNA prepared from *Hydra vulgaris* AEP. PCR amplification was performed using reverse primers containing T7 or SP6 promoter sequences (Qiagen #28506). Amplicons were cloned into the Zero Blunt vector (Invitrogen #K2700-20) and transformed into One Shot™ TOP10 chemically competent *E. coli* (Invitrogen #C404010). Inserts were confirmed by colony PCR and Sanger sequencing. Plasmids were isolated using the Miniprep (Qiagen #27106) kit and served as templates for a second round of PCR to generate transcription templates. Primers are listed in Table 5.

Approximately 250 ng of purified PCR product was used for in vitro transcription with the DIG RNA Labeling Kit (Roche/Sigma #11175025910) and DIG732 U-11 labeling mix (Sigma #11685619910), under RNAse-free conditions using Ambion RNAse-in.RNA probes were purified with the RNA Clean & Concentrator-25 kit (Zymo Research #R1017), diluted to 750 ng in 7.5 μL RNase-free H₂O, and stored at –70°C.

### Fluorescence in Situ Hybridization

A previously published protocol^78^ was used for fluorescence in situ hybridization. Full experimental details are provided in the original publication, and essential steps are summarized below.

*Hydra vulgaris* AEP were washed in HM, relaxed in 2% urethane, and fixed in 4% paraformaldehyde (PFA) for 1 hour at room temperature. Fixed animals were washed in PBST (0.1% Tween-20 in PBS), dehydrated through a methanol series to 100% MeOH, and stored at least overnight at -20°C. Samples were rehydrated, permeabilized with proteinase K (10 µg/mL), quenched with glycine, acetylated in triethanolamine with acetic anhydride, refixed in 4% PFA, and equilibrated in 2X SSC. Prehybridization and hybridization were carried out at 56°C using hybridization solution (50% formamide, 5× SSC, 1× Denhardt’s, 100 µg/mL heparin, 0.1% Tween-20, 0.1% CHAPS). DIG- and/or FITC-labeled RNA probes (∼180 ng) were prepared as described above and hybridized for ∼65 hours.

Post-hybridization washes were performed at 56°C with decreasing HS/SSC concentrations, followed by washes in 2× SSC + 0.1% CHAPS and MABT. Samples were blocked in MABT + 1% BSA and 20% sheep serum, then incubated overnight at 4°C with anti-DIG-POD (Sigma #11207733910, 1:2000). Signal was developed by tyramide amplification using Alexa Fluor 488 or 594 tyramide (1:100) prepared in borate buffer with dextran sulfate, H_2_O_2_, and 4-iodophenol. After quenching in glycine, tissues were washed, counterstained with Hoechst (1:1000), and dehydrated through a glycerol series before mounting in 80% glycerol with a spacer (EMS #703279S).

### Fluorescence image acquisition

Fluorescent images of FISH and immunofluorescence experiments were acquired using a Zeiss 980 laser scanning microscope with a 10X Plan-Apochromat objective (10x/0.45), Images were collected at 1024×1024 resolution with a pinhole size of 1AU and a Z-step size of 3.3µm.

Imaging parameters were set using ZEN 568 software using two tracks, one to image Hoechst and one to image the FISH or immunofluorescence signal. Whole animals and regenerates were imaged by tiling multiple Z-stacks. Maximum intensity projections were generated in Zeiss Zen Blue 3.7.

### Fluorescence in Situ Hybridization analysis

Fluorescence intensity ratios at the wound edge were quantified by measuring the intensity profile along a 100µm-wide line drawn from the wound tip into the body column. The average intensity from 0-200 µm was divided by the average intensity from 300-500µm, and this ratio was normalized to body column width. Area of the FISH signal was measured by manually outlining the expression domain and normalizing the area to body column width.

### p-ERK immunofluorescence analysis

Wound p-ERK fluorescence signal was quantified by plotting intensity profile along a line drawn from the wound tip towards the body column. The average intensity in the first 100 µm was used to calculate the proximal p-ERK fluorescence intensity, and the average intensity 200-400µm from the wound edge was used to calculate the distal intensity. Wound edge p-ERK enrichment after DMSO or U0126 treatment was calculated as the ratio of proximal to distal intensity. As a control, we confirmed that wound-edge p-ERK signal does not reflect nonspecific antibody binding at the cut surface (Supp. Fig. 3B).

### pH3 immunofluorescence analysis

Nuclei stained with Hoechst that are positive for phospho-histone-H3 were manually counted by examining each maximum projection.

## SUPPLEMENTAL FIGURES

**Supp. Fig. 1.**
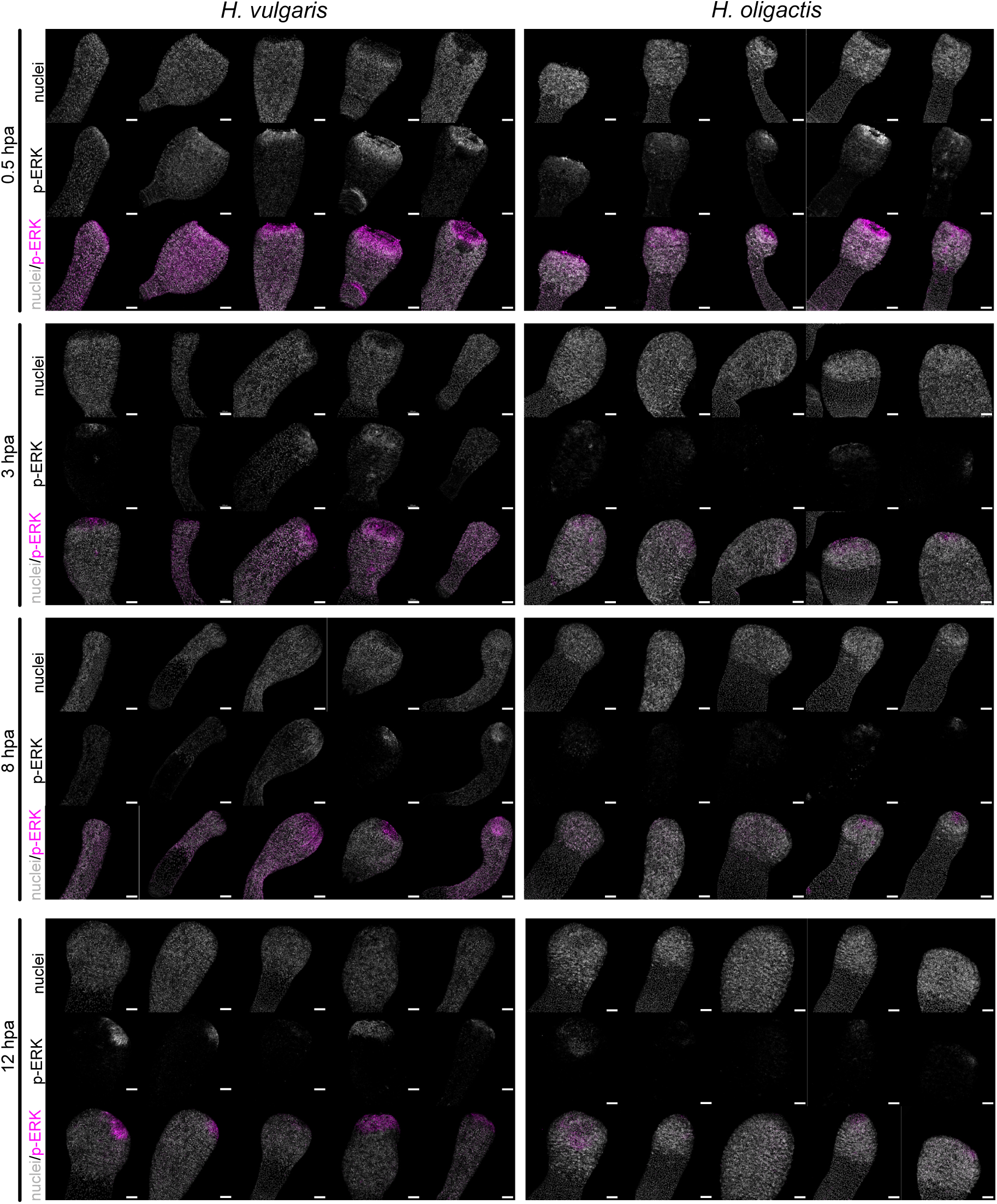
Additional p-ERK immunofluorescence images of regenerating *Hydra.* Immunofluorescence images showing p-ERK and Hoechst staining in regenerating heads of *H. vulgaris* and *H. oligactis* at 0.5, 3, 8, and 12 hpa to accompany Figures 1 and 7. Scale bar = 100μm.

**Supplemental Fig. 2.**
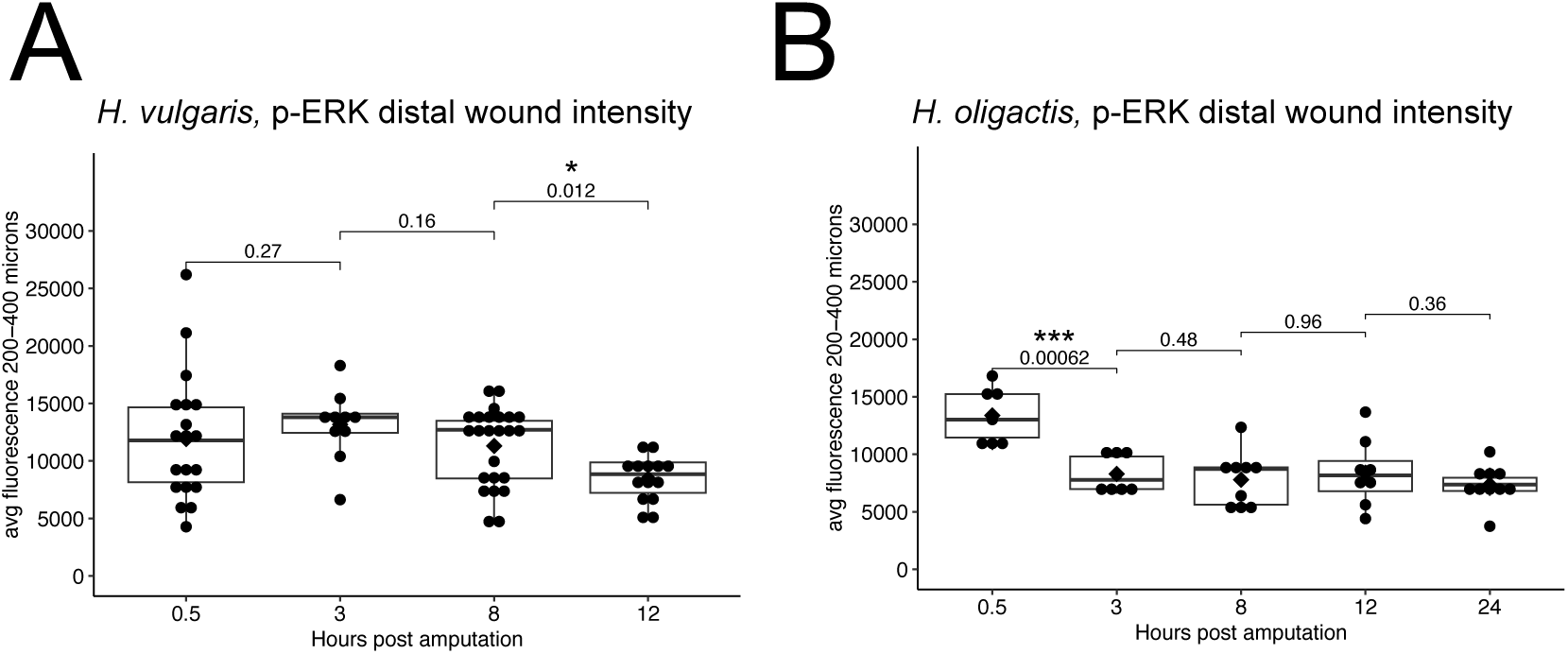
*H. vulgaris* demonstrates a prolonged persistence of p-ERK at the distal wound as compared to *H. oligactis*. (A) Quantification of the distal p-ERK signal intensity in regenerating *H. vulgaris* heads corresponding to Figure 1C. Each point represents an individual animal (0.5 hpa n=19, 3 hpa n=10, 8 hpa n=24, 12 hpa n=14). Fluorescence intensity was quantified using a plot profile analysis, statistical comparisons were performed using the Wilcoxon rank-sum test. (B) Quantification of the distal p-ERK signal intensity in regenerating *H. oligactis* heads corresponding to Figure 7C. Each point represents an individual animal (0.5hpa n=7, 3hpa n=8, 8 hpa n=9, 12 hpa n=8, 24 hpa n=11). Fluorescence intensity was quantified using a plot profile analysis, statistical comparisons were performed using the Wilcoxon rank-sum test.

**Supp. Fig. 3.**
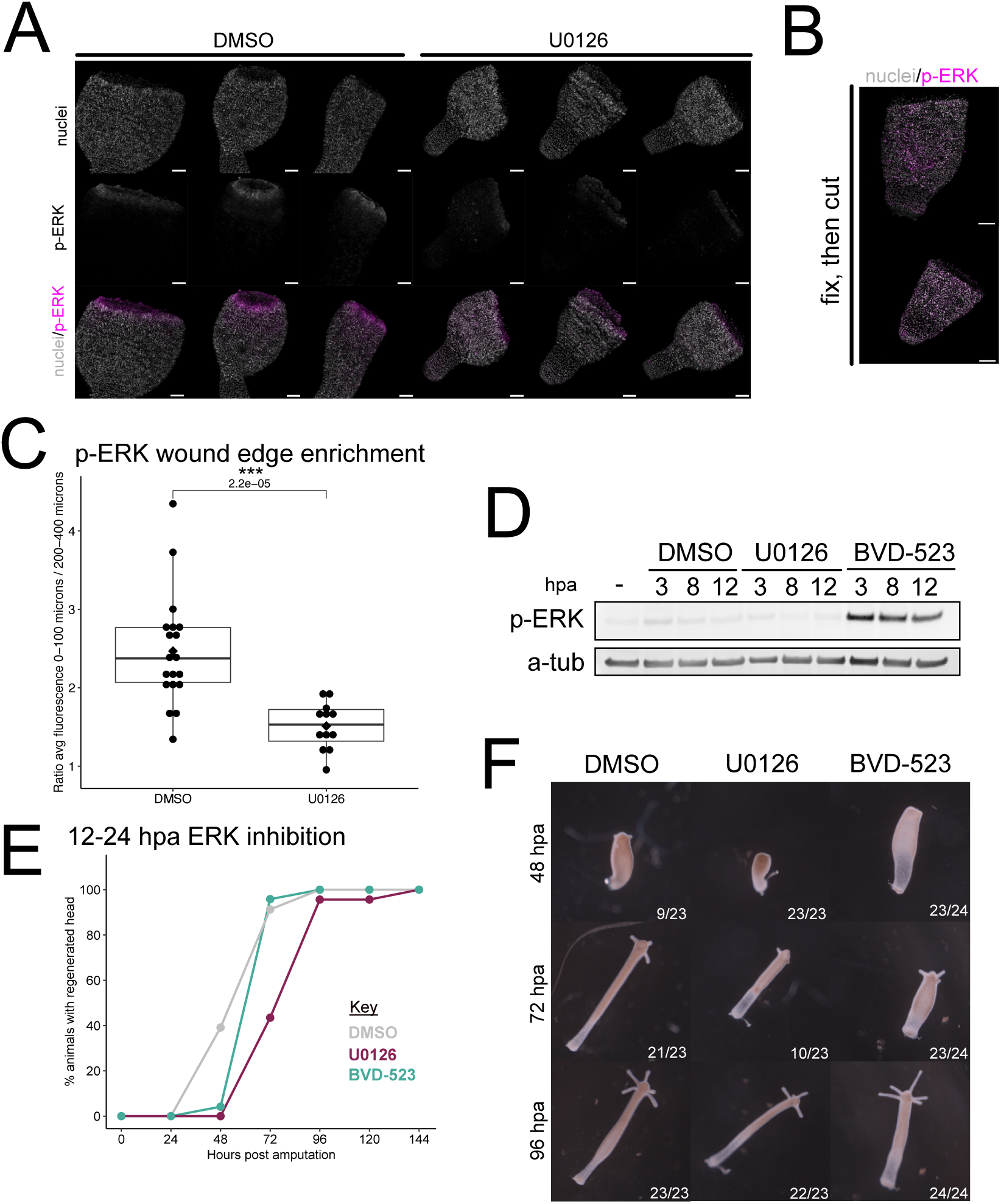
p-ERK dynamics and requirement for head regeneration in *H. vulgaris.* (A) Immunofluorescence images of regenerating *H. vulgaris* heads at 0.5 hpa treated with DMSO or U0126, stained for p-ERK together and Hoechst to label nuclei. Scale = 100µm. (B) Immunofluorescence images of regenerating *H. vulgaris* heads that were bisected after fixation and stained for p-ERK and Hoechst to label nuclei. Scale = 100µm. (C) Quantification of p-ERK signal enrichment at the wound edge from panel A, with each point representing an individual animal (DMSO n=19, U0126, n=12). Statistical comparisons were performed using the Wilcoxon rank-sum test. (D) Western Blot analysis of p-ERK and α-tubulin levels in regenerating *H. vulgaris* heads. Animals not treated with U0126 or BVD-523 received DMSO as a vehicle control. The first lane shows whole, uninjured animals without treatment, indicating basal levels of p-ERK. (E) Head regeneration rates for *H. vulgaris* treated with ERK inhibitors (U0126, 25uM, burgundy; BVD-523, 7.5uM, teal) or a vehicle control (DMSO, gray) for 12-24 hpa, followed by washout into *Hydra* Medium for the remainder of the experiment. (DMSO n=21, BVD-523 n=24, U0126 n=23). (F) Representative images of *H. vulgaris* head regeneration at 48, 72, 96 hpa from treatments shown in panel E.

**Supp. Fig. 4.**
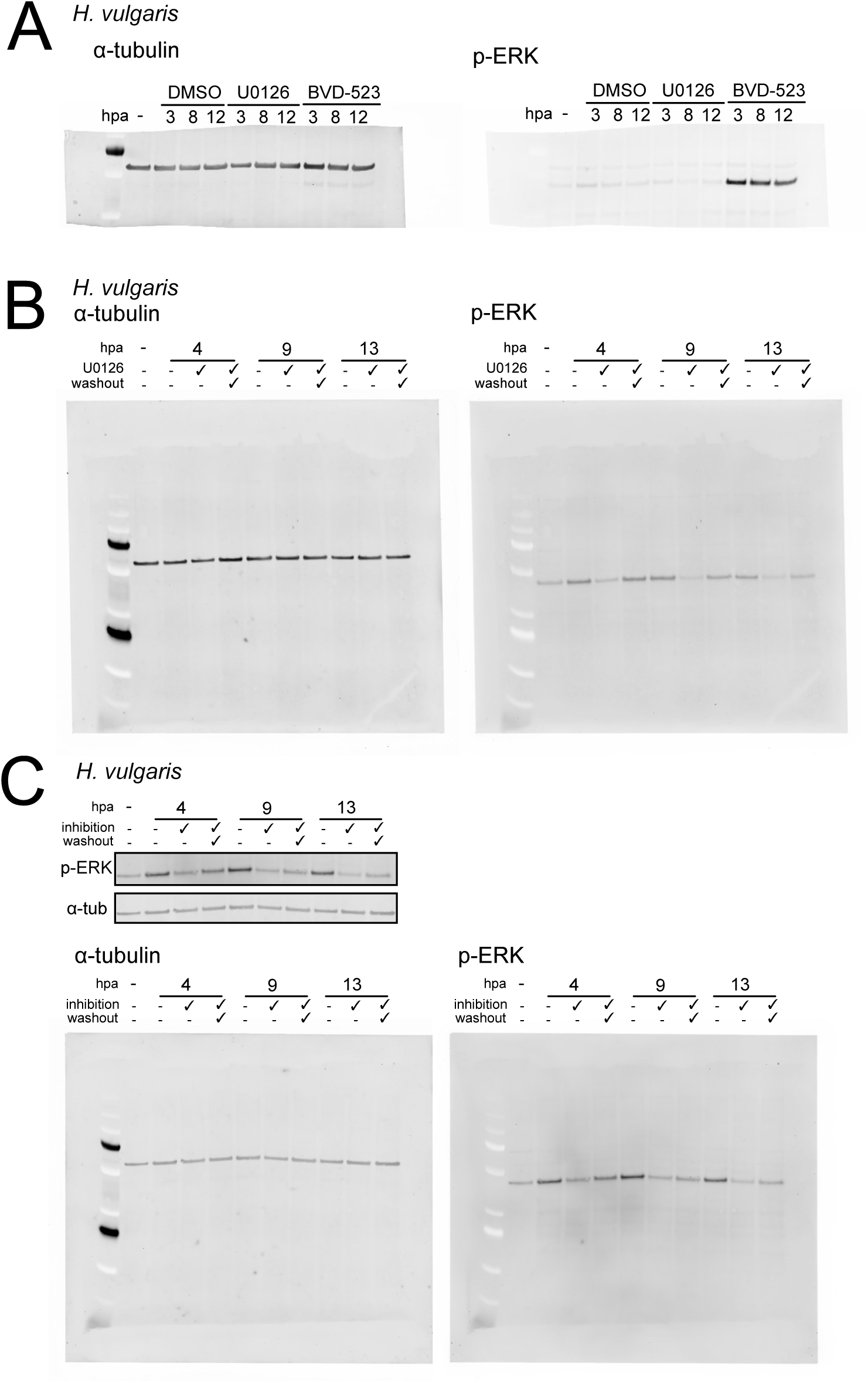
Uncropped Western blot data and biological replicate. (A) Uncropped Western blot corresponding to Supplemental Figure 2C. (B) Uncropped Western blot corresponding to Figure 3B. (C) Biological replicate for experiments shown in Figure 3B. Both the cropped and uncropped versions are shown.

**Supp. Fig. 5.**
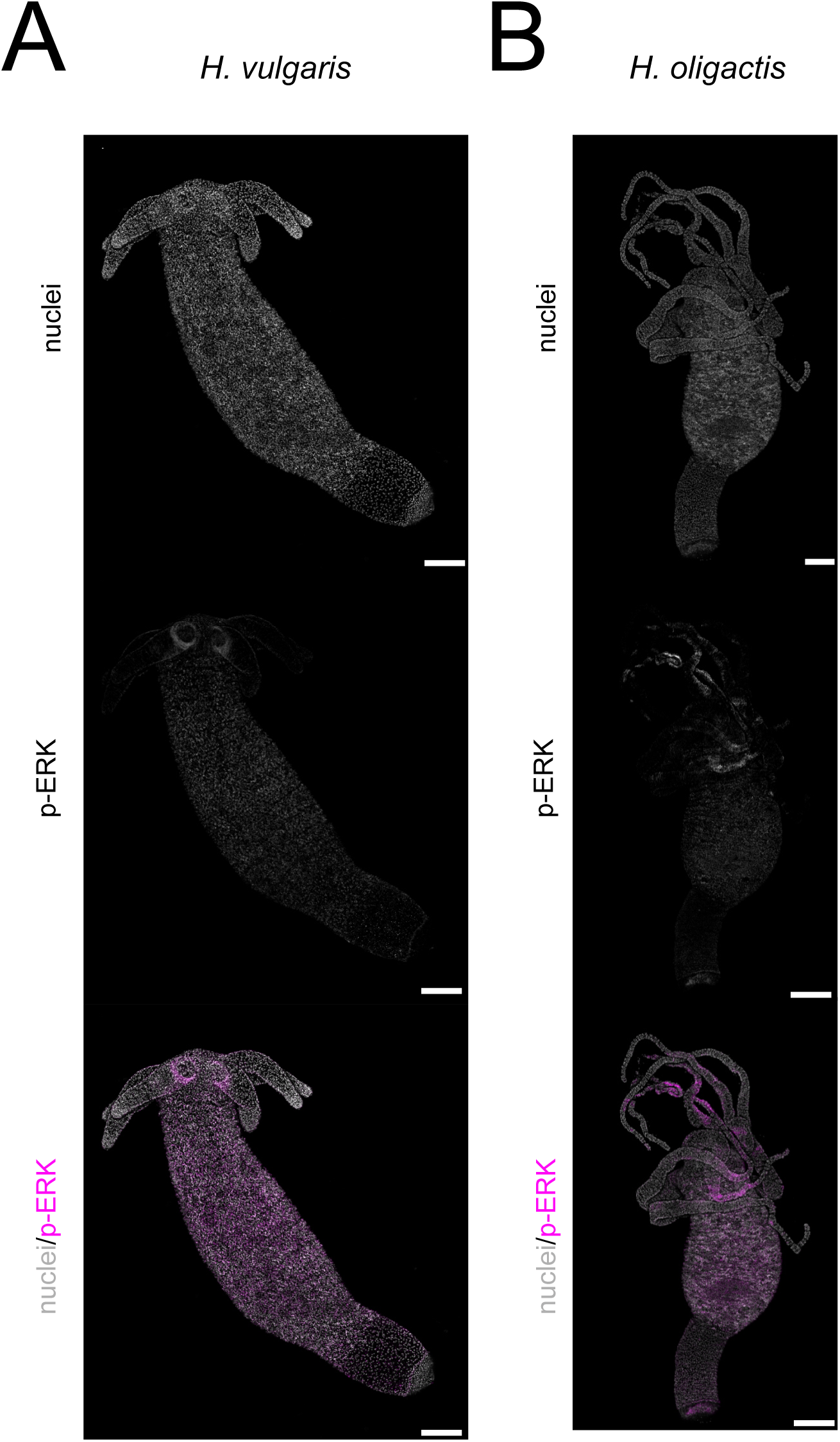
Basal levels of p-ERK in uninjured *H. vulgaris* and *H. oligactis*. (A) Immunofluorescence images of uninjured *H. vulgaris* stained for p-ERK and Hoechst to label nuclei, showing detectable basal p-ERK expression. Scale bar = 200µm. (B) Immunofluorescence images of uninjured *H. oligactis* using p-ERK and Hoechst to label nuclei, showing detectable expression. Scale bar = 200µm.

**Supp. Fig. 6.**
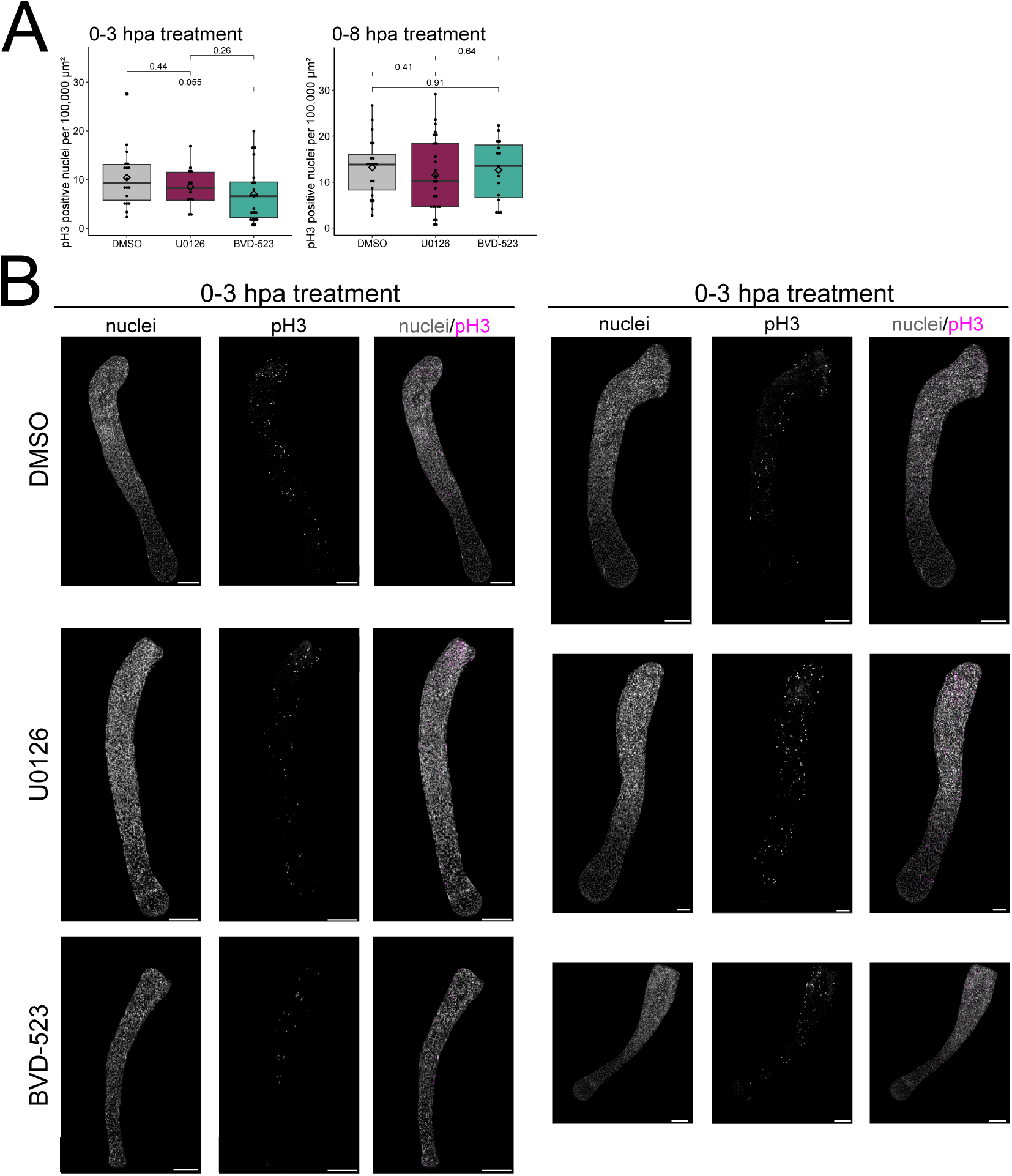
Transient ERK inhibition during the first 3 or 8 hours of head regeneration does not significantly alter proliferation. (A) Quantification of pH3-positive nuclei normalized to tissue area in animals treated with ERK inhibitors during the first 3 or 8 hpa. Each dot represents an individual animal. For 0–8 hpa treatments, sample sizes were DMSO n = 20, U0126 n = 26, and BVD-523 n = 15. For 0–3 hpa treatments, sample sizes were DMSO n = 18, U0126 n = 12, and BVD-523 n = 22. Statistical comparisons were performed using the Wilcoxon rank-sum test. (B) Representative immunofluorescence images of regenerating animals stained for pH3 and Hoechst following ERK inhibitor treatment during the first 3 or 8 hours of regeneration. Scale bar = 200 µm.

**Supp. Fig. 7.**
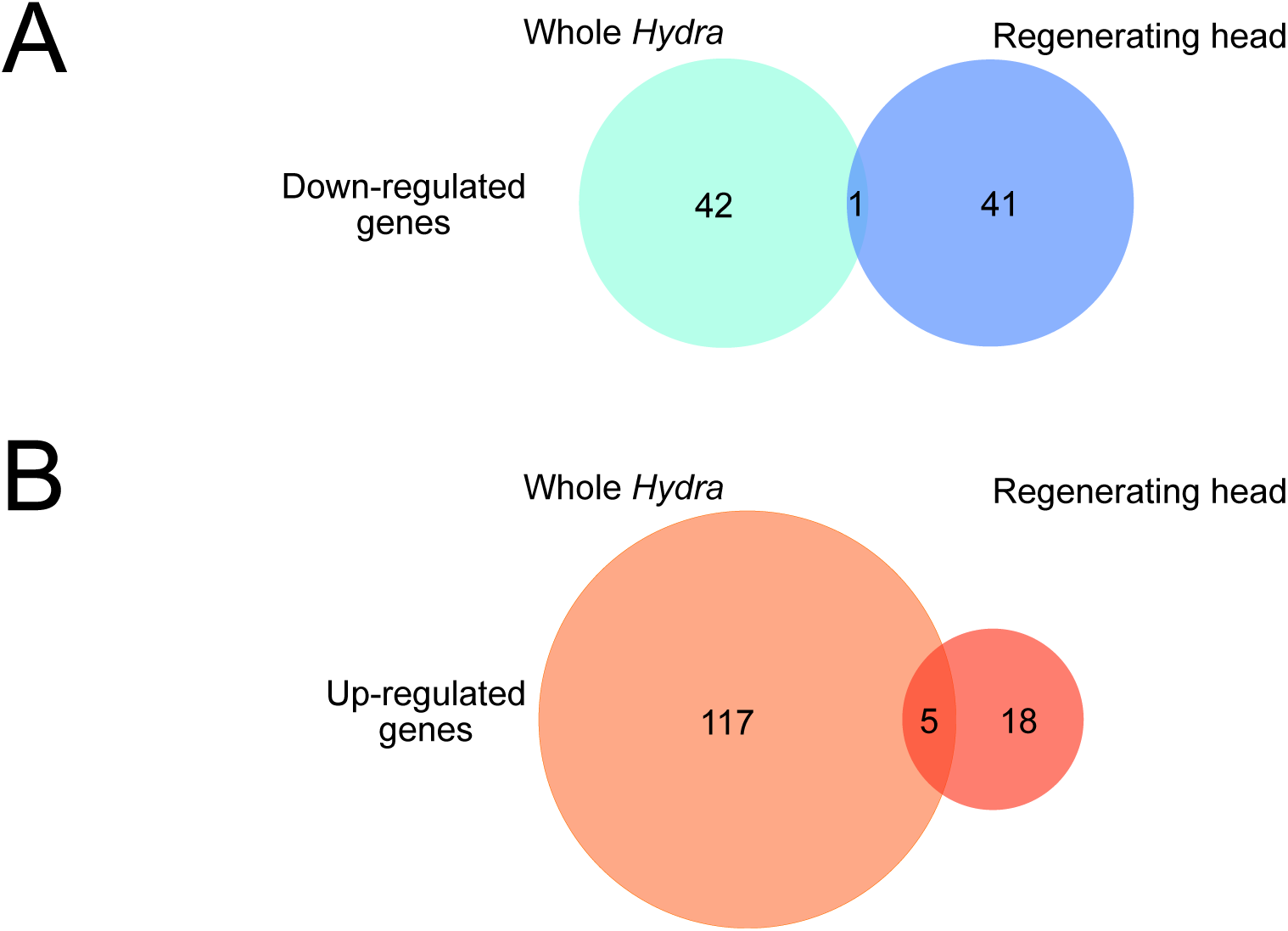
Distinct differentially expressed genes (DEGs) in whole animals and regenerating heads following U0126 treatment. (A) Venn diagram showing downregulated DEGs identified in whole animals and regenerating heads. The complete list of genes is provided in Supplemental Table 2. (B) Venn diagram showing upregulated DEGs identified in whole animals and regenerating heads. The complete list of genes is provided in Supplemental Table 2.

**Supp. Fig. 8.**
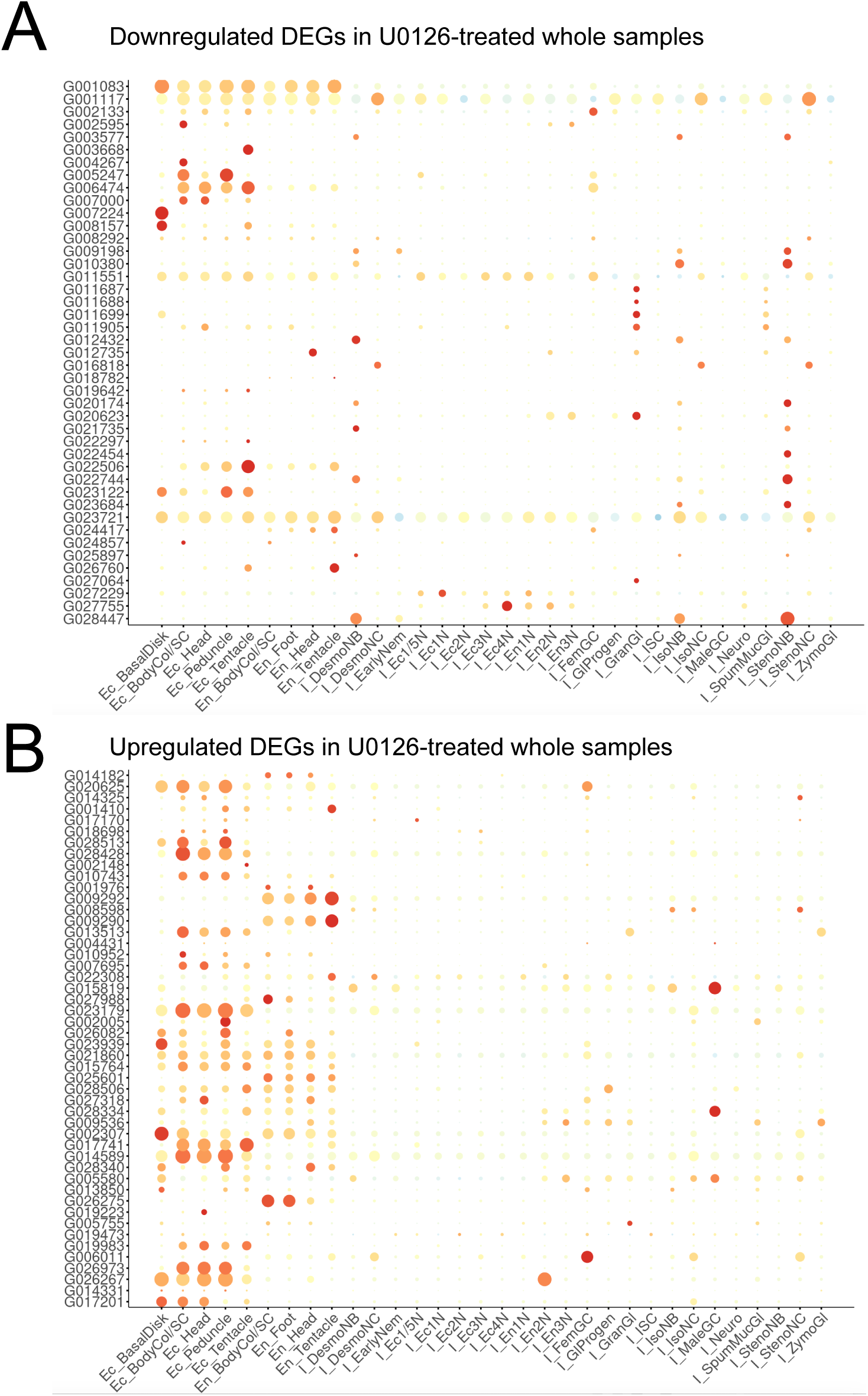
Differentially expressed genes (DEGs) in whole animals following U0126 treatment are predominantly expressed in epithelial cells. Dot plots of previously published single-cell RNA-seq data showing expression of selected DEGs identified in uninjured animals after U0126 treatment. The y-axis indicates gene identifiers, and the x-axis indicates cell types. (A) Genes downregulated following U0126 treatment. (B) Genes upregulated following U0126 treatment.

**Supp. Fig. 9.**
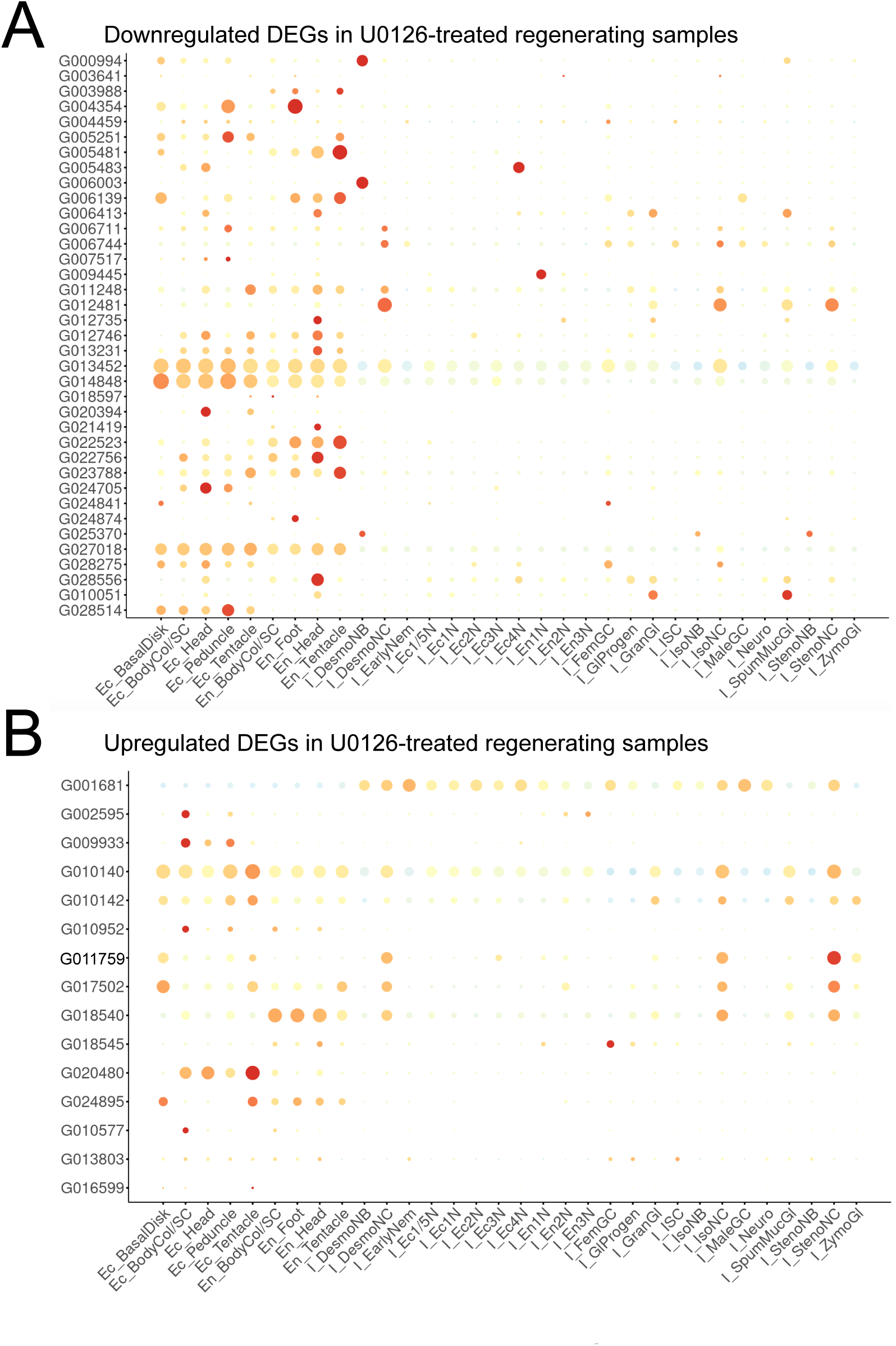
Differentially expressed genes (DEGs) in regenerating heads following U0126 treatment are predominantly expressed in epithelial cells. Dot plots of previously published single-cell RNA-seq data showing expression of selected DEGs identified in regenerating heads after U0126 treatment. The y-axis indicates gene identifiers, and the x-axis indicates cell types. (A) Genes downregulated following U0126 treatment. (B) Genes upregulated following U0126 treatment.

**Supp. Fig. 10.**
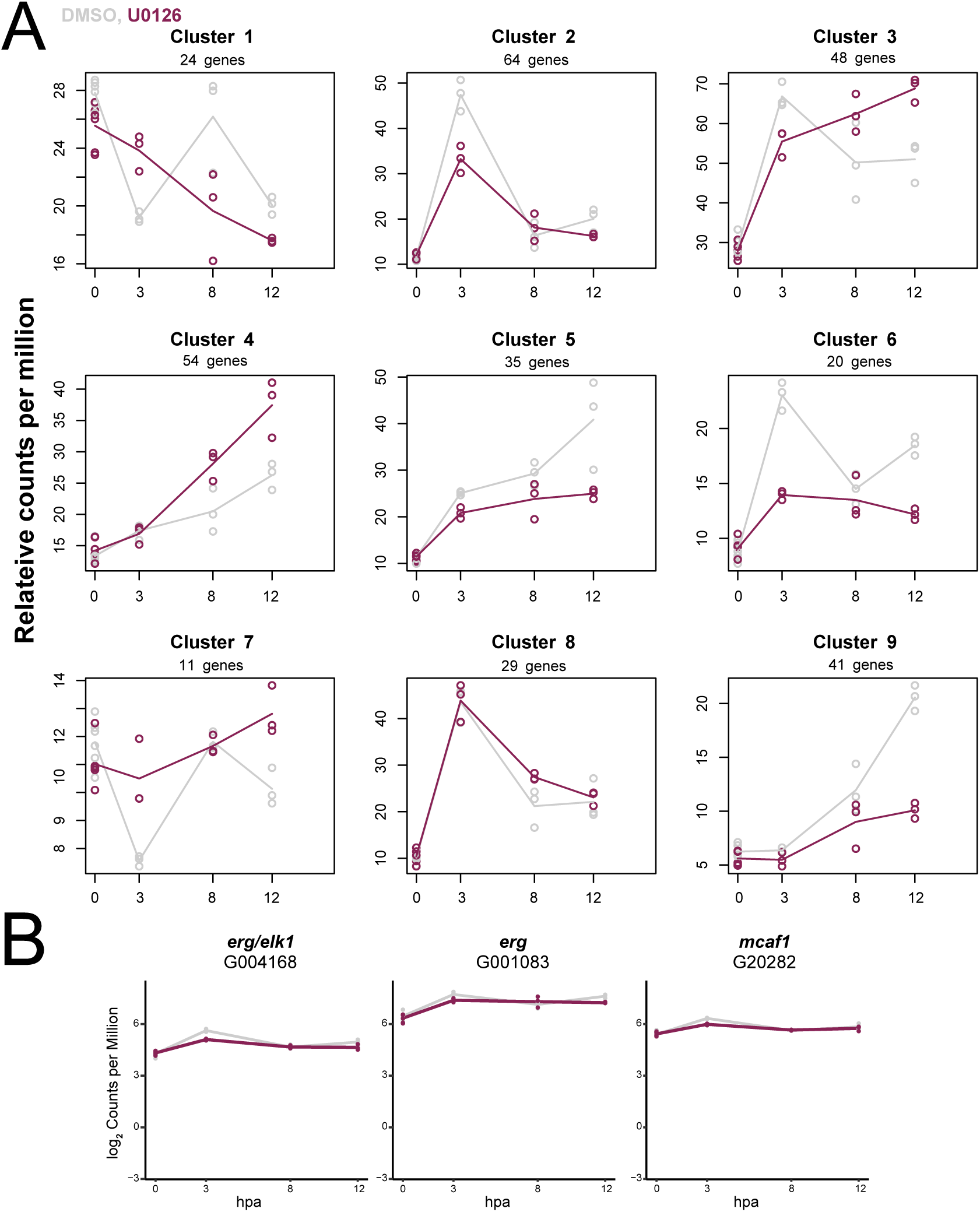
ERK signaling influences the transcription of genes activated during early and late head regeneration. (A) Clustering of RNA-seq data using maSigPro analysis to identify temporally regulated gene expression patterns. The full gene list is provided in Supplemental Table 3. (B) Expression levels of ETS family transcription factors (G004168 and G001083) and *mcaf1* transcripts. Gene names correspond to annotations from the *Hydra* AEP Genome Project.

**Supp. Fig. 11.**
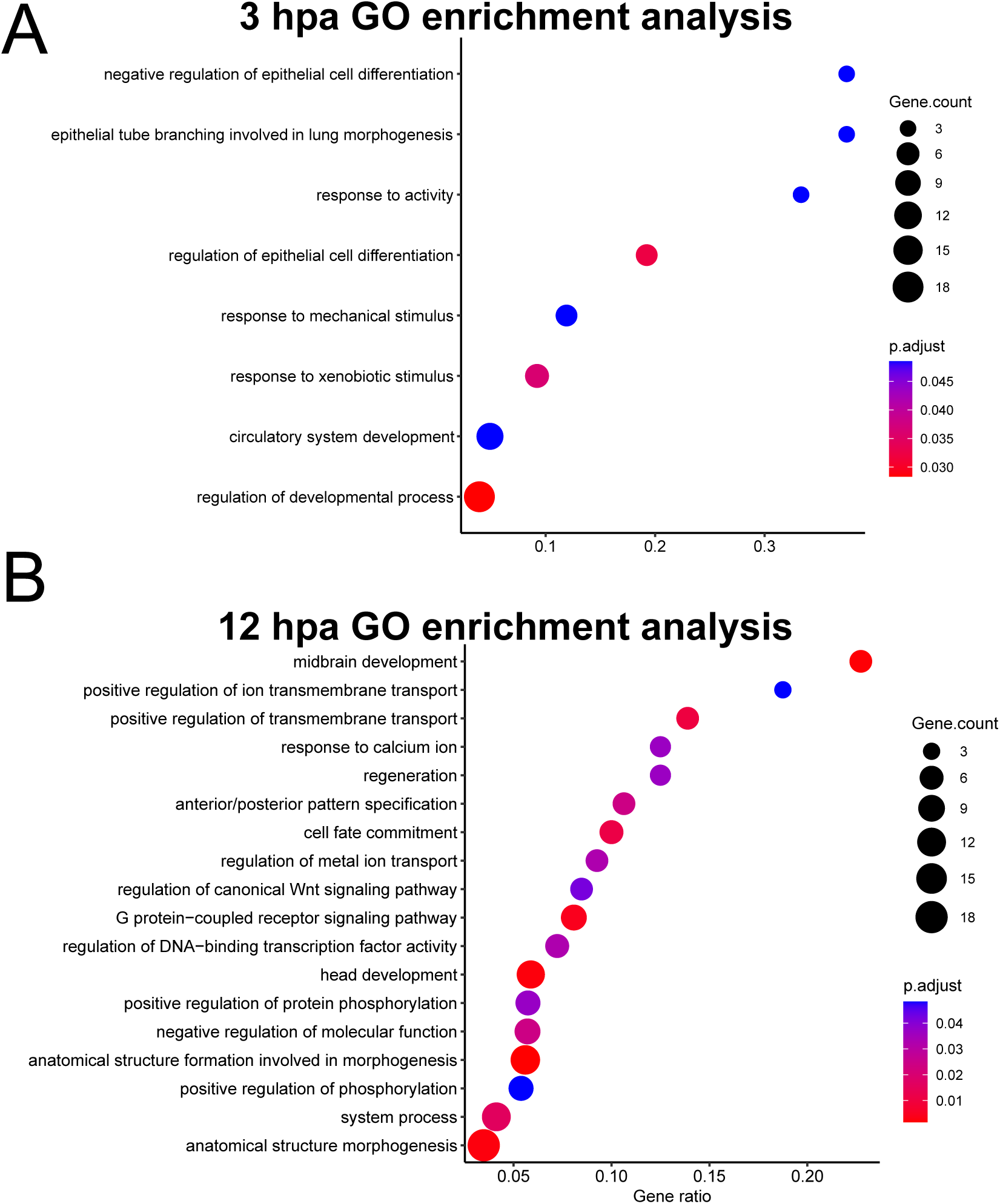
Gene Ontology (GO) term analysis reveals increased enrichment of morphological and patterning-related categories at 12 hours post-amputation. (A) GO term analysis comparing transcriptional trends between U0126- and DMSO-treated samples at 3 hpa. (B) GO term analysis comparing transcriptional trends between U0126- and DMSO-treated samples at 12 hpa.

**Supp. Fig. 12.**
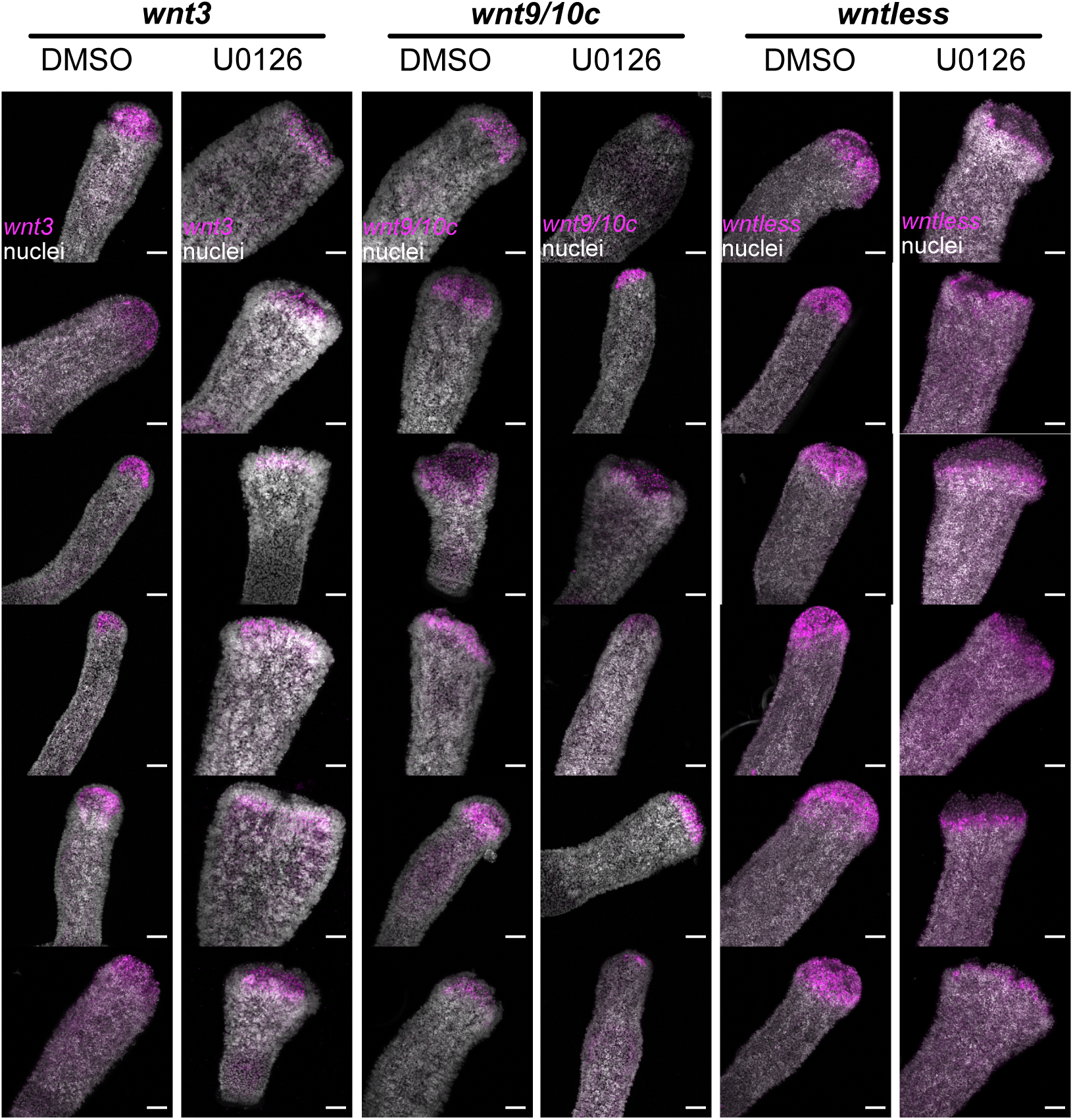
Additional fluorescence in situ hybridization (FISH) images of Wnt gene expression accompanying Fig. 6A. Representative images of regenerating head tips treated with DMSO or U0126 during the first 12 hours of regeneration and fixed at 12 hpa. Animals were stained by FISH (magenta) and counterstained with Hoechst (white). Scale bar = 100 µm.

**Supp. Fig. 13.**
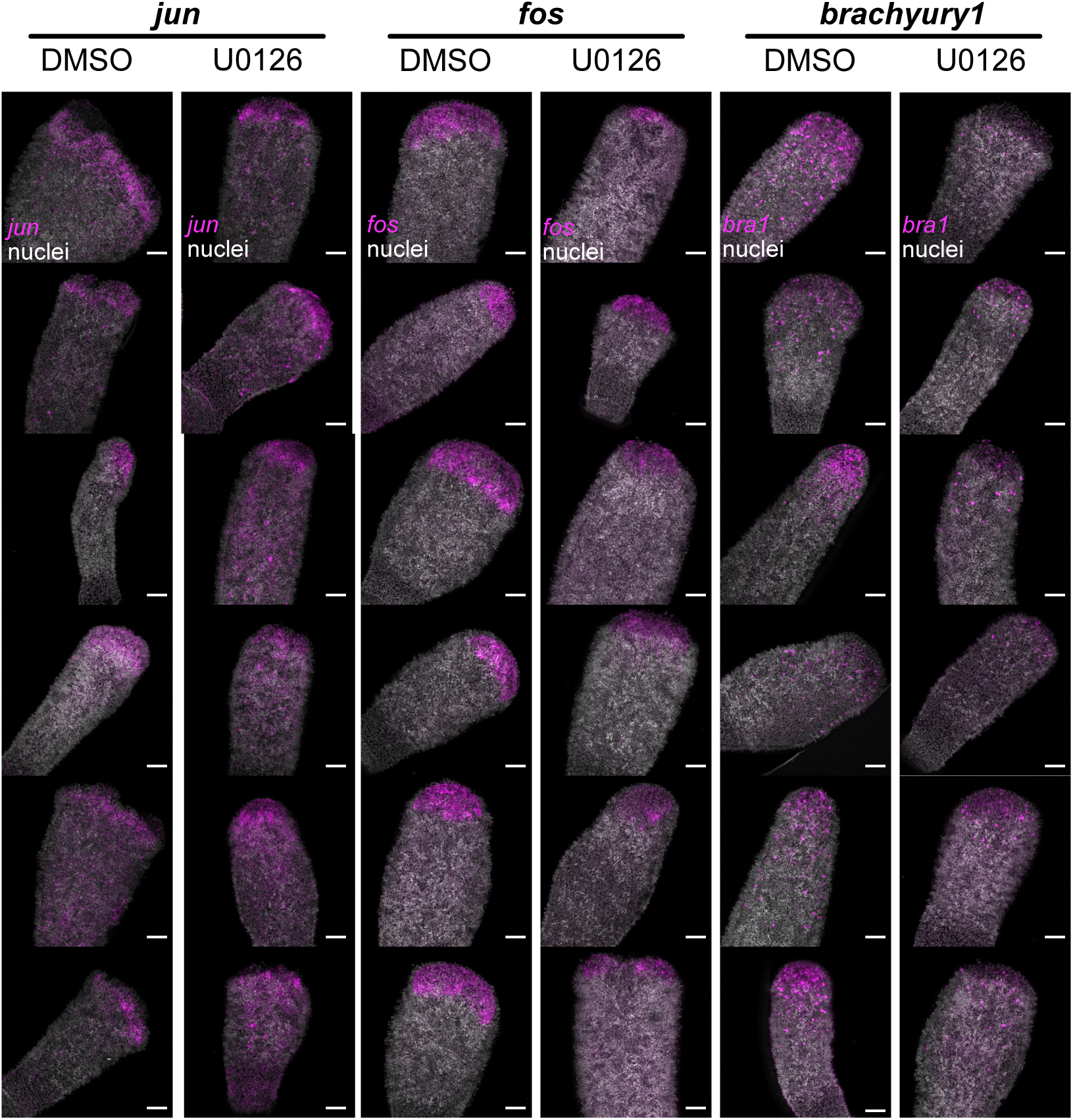
Additional fluorescence in situ hybridization (FISH) images of jun, fos, and bra1 expression accompanying Fig. 6C,D. Representative images of regenerating head tips treated with DMSO or U0126 during the first 12 hours of regeneration and fixed at 12 hpa. Animals were stained by FISH (magenta) and counterstained with Hoechst (white). Scale bar = 100 µm.

**Supp. Fig. 14.**
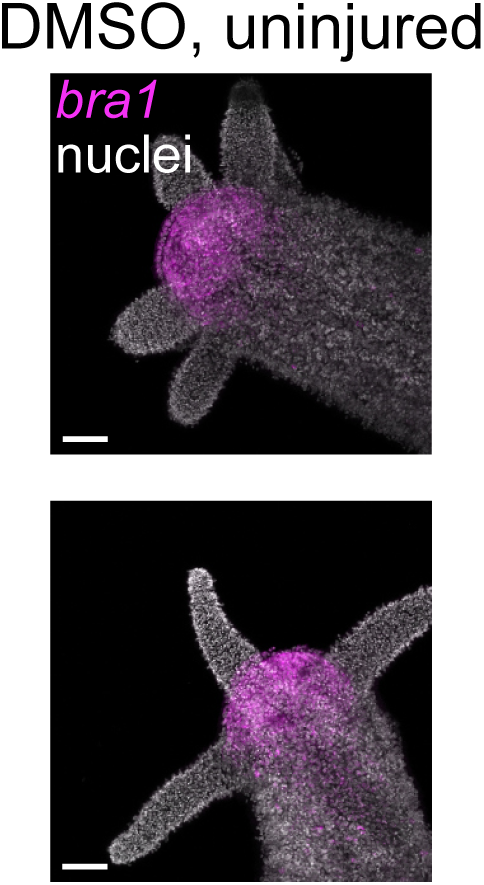
Brachyury1 expression in the head of an uninjured *H. vulgaris*. Representative images of uninjured animals treated with DMSO showing *brachyury1* (*bra1*) expression in the head. Animals were stained by fluorescence in situ hybridization (FISH, magenta) and counterstained with Hoechst (white). Scale bar = 100 µm.

**Supp. Fig. 15.**
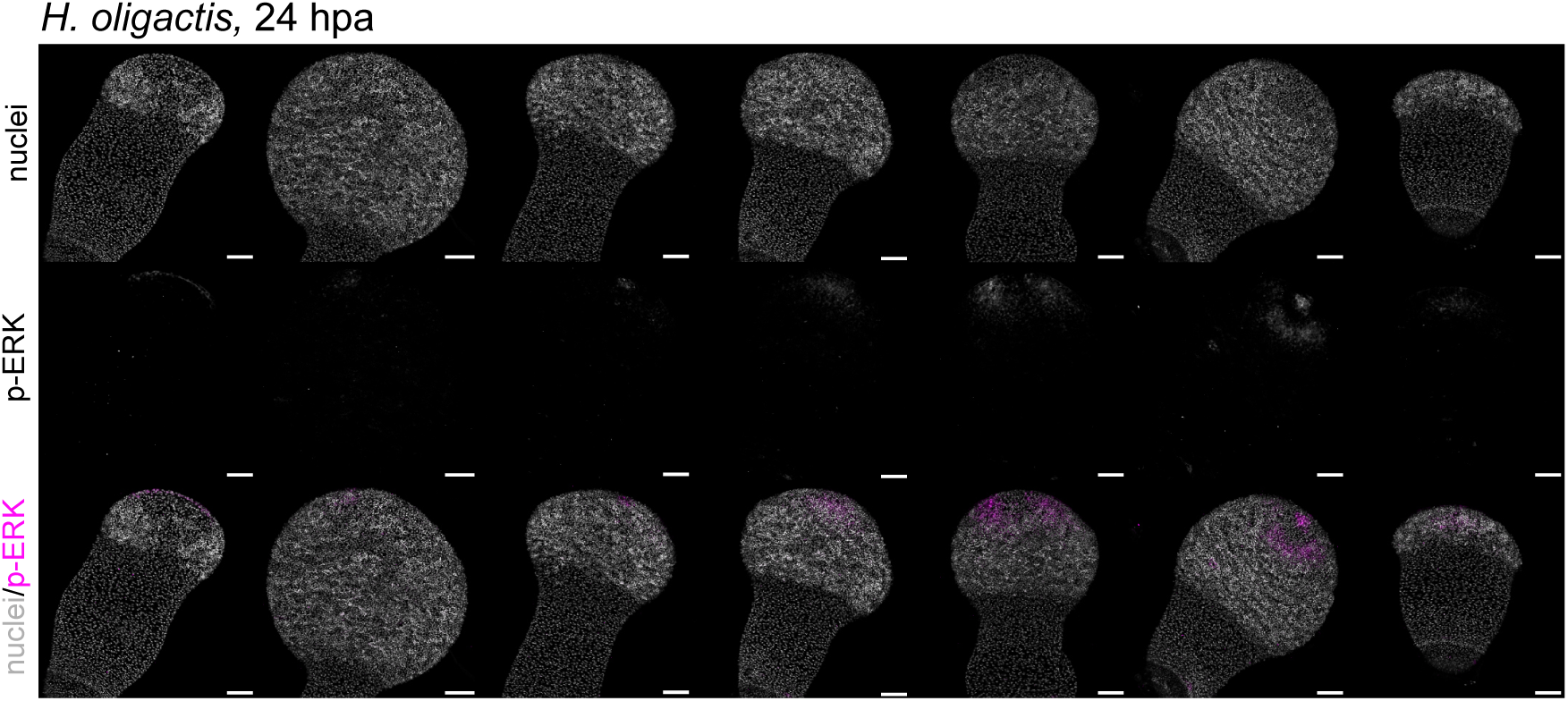
Additional immunofluorescence images accompanying Fig. 7 showing ERK activation in *H. oligactis* at 24 hpa. Representative images of regenerating *H. oligactis* heads stained for phosphorylated ERK (p-ERK) and counterstained with Hoechst. Scale bar = 100 µm.

**Supp. Fig. 16.**
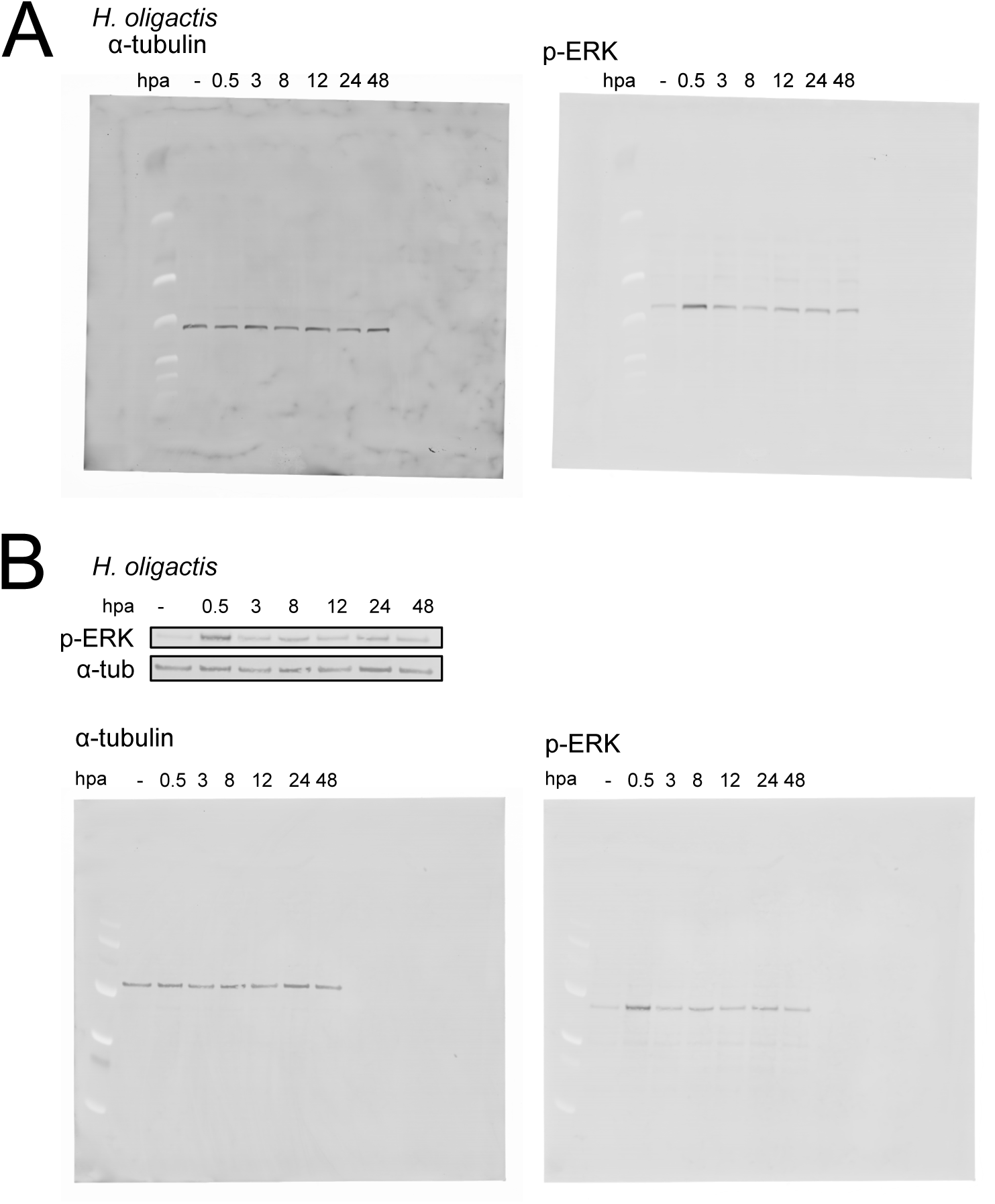
Uncropped Western blot images accompanying Fig. 7C and biological replicate. (A) Uncropped Western blot corresponding to the experiment shown in Fig. 7C. (B) Biological replicate for experiments shown in Fig. 7C. Both the cropped and uncropped versions are shown.

**Supp. Fig. 17.**
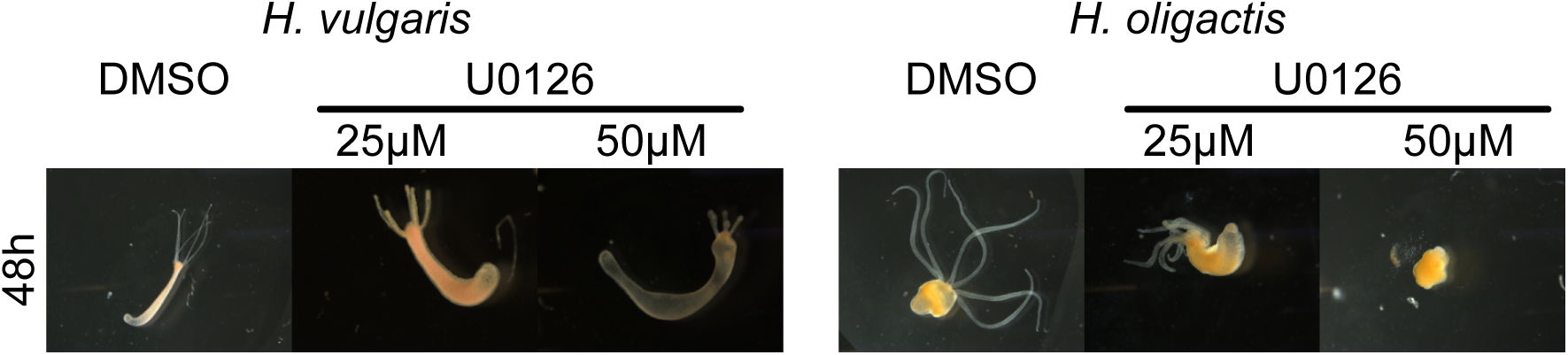
Effects of DMSO and U0126 on uninjured *H. vulgaris* and *H. oligactis*. Representative images of uninjured *H. vulgaris* and *H. oligactis* treated with DMSO (vehicle control) or U0126 at 25 µM or 50 µM for 48 hours. Treatment with 50 µM U0126 resulted in cell sloughing and was therefore not used in subsequent experiments.

**Supp. Fig. 18.**
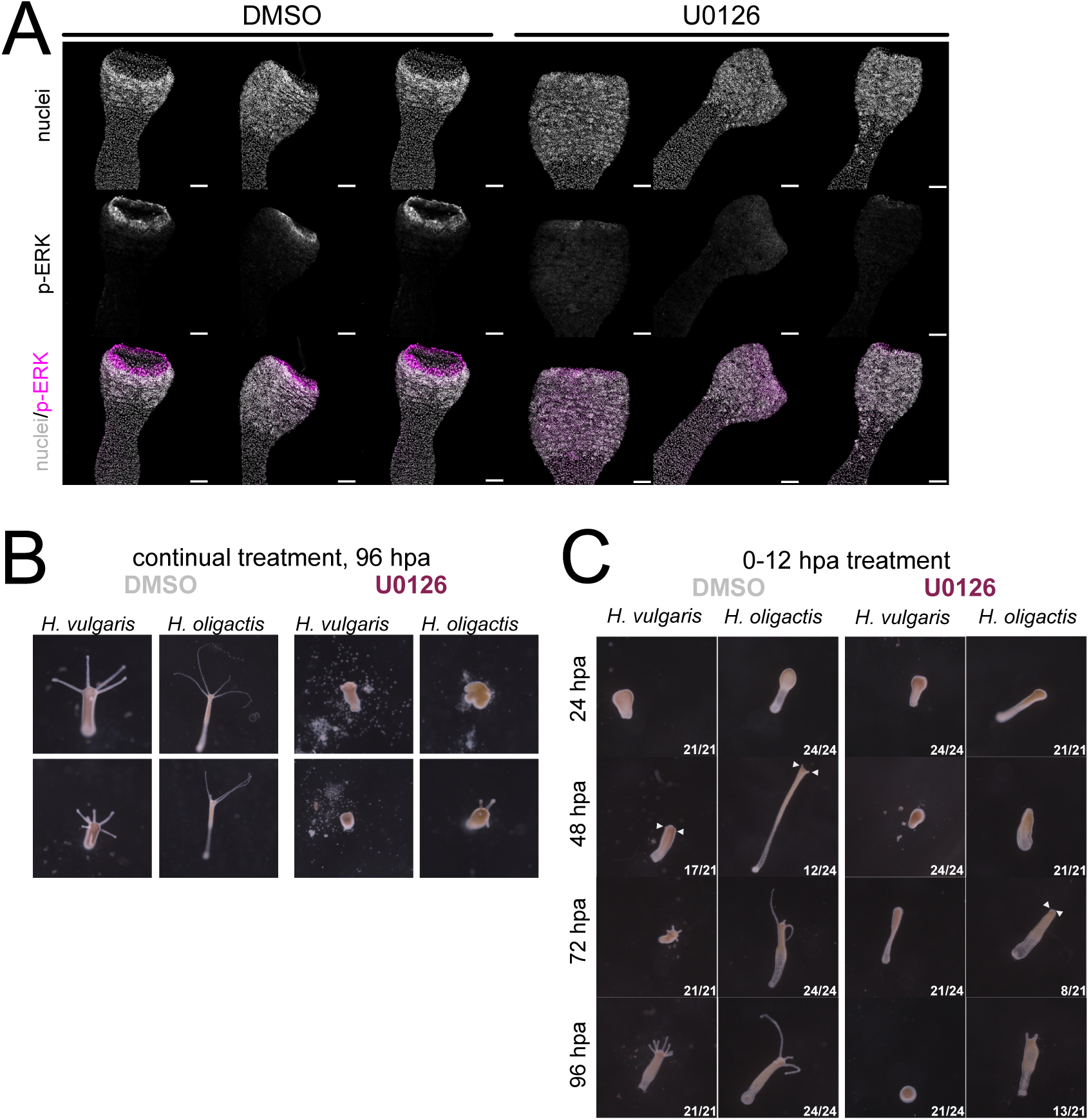
Additional analyses of the effects of DMSO and U0126 on Hydra vulgaris and Hydra oligactis. (A) Additional immunofluorescence images accompanying Fig. 8A showing p-ERK and Hoechst staining in regenerating *H. oligactis* heads at 0.5 hours hpa following treatment with DMSO or U0126. Scale bar = 100 µm. (B) Representative images accompanying Fig. 8C of regenerating *H. vulgaris* and *H. oligactis* continuously treated with DMSO or U0126 and imaged at 96 hpa. (C) Representative images accompanying Fig. 8D of *H. vulgaris* and *H. oligactis* treated with DMSO or U0126 from 0–12 hpa.

## SUPPLEMENTAL TABLES

Supp.Table 1. **RNA-seq contrast tables listing upregulated and downregulated differentially expressed genes (DEGs) from the regenerating head dataset.** Gene IDs correspond to annotations from the *Hydra vulgaris* AEP Project Portal.^73^

Supp. Table 2. **Differentially expressed genes (DEGs) shared between, and unique to, the whole-animal and regenerating head RNA-seq datasets.** Gene IDs correspond to annotations from the *Hydra vulgaris* AEP Project Portal.^73^

Supp.Table 3. **Cluster analysis of RNA-seq data.** List of all genes and their assignment to clusters identified through maSigPro analysis. Gene IDs correspond to annotations from the *Hydra vulgaris* AEP Project Portal.

**Supp.Table 4.**
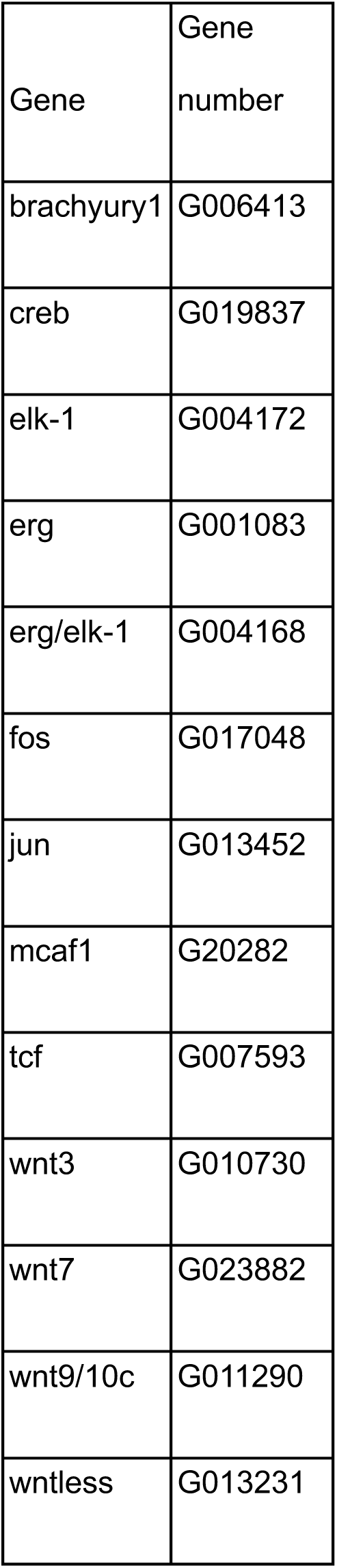
Gene reference table listing all genes referenced in this study and their corresponding identifiers. Gene IDs correspond to annotations from the *Hydra vulgaris* AEP Project Portal (https://research.nhgri.nih.gov/HydraAEP/).^73^

| Gene | Gene number |
| --- | --- |
| brachyury1 | G006413 |
| creb | G019837 |
| elk-1 | G004172 |
| erg | G001083 |
| erg/elk-1 | G004168 |
| fos | G017048 |
| jun | G013452 |
| mcaf1 | G20282 |
| tcf | G007593 |
| wnt3 | G010730 |
| wnt7 | G023882 |
| wnt9/10c | G011290 |
| wntless | G013231 |

**Supp. Table 5.**
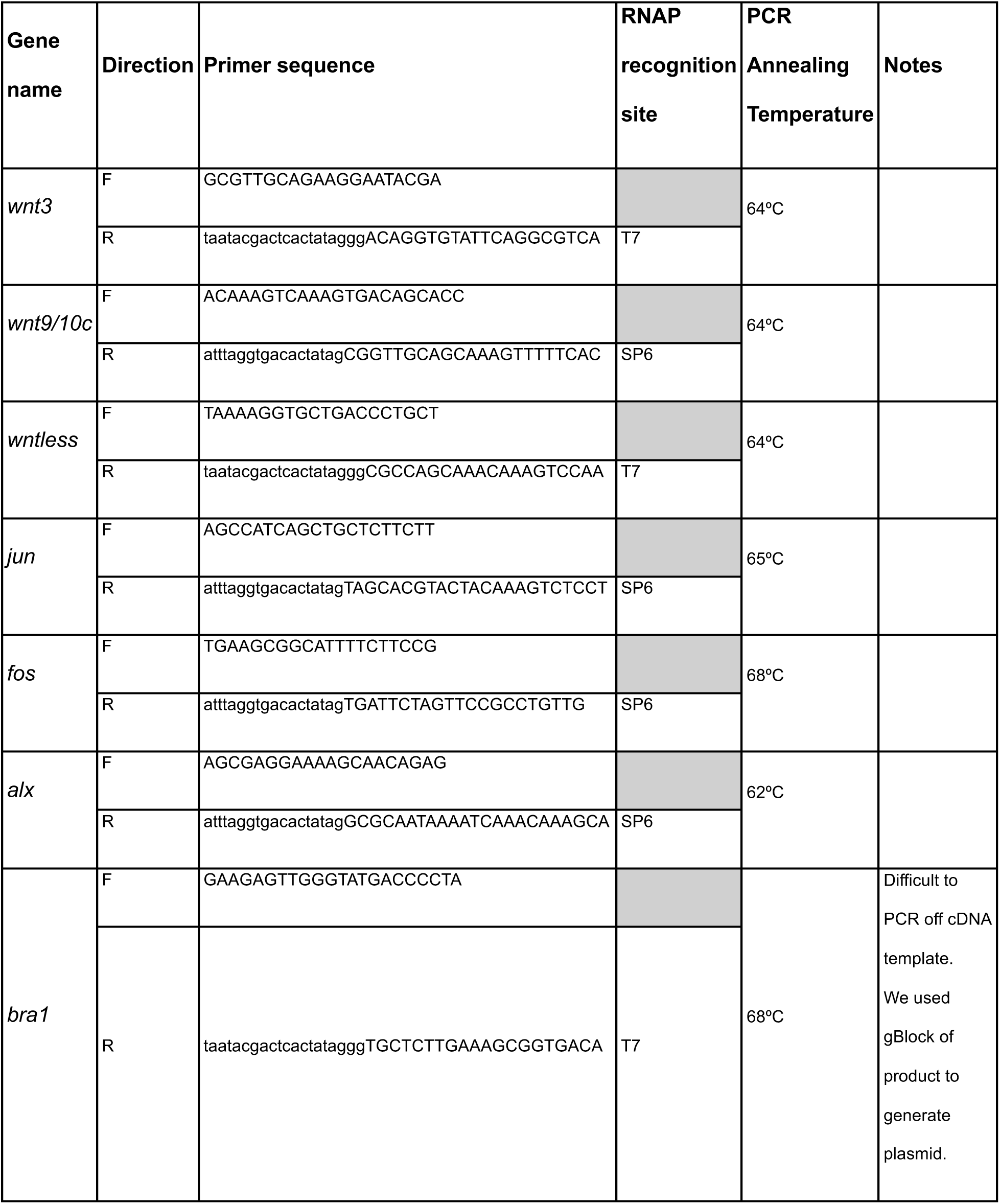
Primer sequences used for fluorescence in situ hybridization (FISH) experiments in this study. Lowercase letters in the reverse primer sequences indicate the RNA polymerase binding site.

## ACKNOWLEDGEMENTS

We would like to thank Dr. Crystal Rogers, Dr. John Albeck, Dr. Bruce Draper, Dr. Alice Accorsi, and members of the Juliano lab for their input and feedback.

This work was supported by the National Institutes of Health grant 5R35GM133689 to CEJ.

This study was made possible in part through access to the MCB Light Microscopy Core Facility, the NIH grant S10OD026702 that allowed funding of the Zeiss 980 LSM and training and support by Thomas Wilkop.

